# Impact of aging on the GABA_B_ receptor-mediated connectome

**DOI:** 10.1101/2024.07.31.606013

**Authors:** Elena Vásquez, Ana L. López, Gerardo M. Oresti, María D. Paez, Eduardo A. Callegari, Diego Masone, Estela M. Muñoz

**Affiliations:** Instituto de Histología y Embriología de Mendoza (IHEM), Universidad Nacional de Cuyo, Consejo Nacional de Investigaciones Científicas y Técnicas (CONICET), Mendoza, Argentina; Instituto de Investigaciones Bioquímicas de Bahía Blanca, Consejo Nacional de Investigaciones Científicas y Técnicas (CONICET) y Departamento de Biología, Bioquímica y Farmacia, Universidad Nacional del Sur (UNS), Bahía Blanca, Argentina; Division of Basic Biomedical Sciences, University of South Dakota Sanford School of Medicine, Vermillion, SD, USA

**Keywords:** GABA_B_R, KCC2, cholesterol, aging, cerebellum

## Abstract

GABA B receptors (GABA_B_Rs) are heterodimeric seven-transmembrane receptors that interact with a range of proteins and form large protein complexes on cholesterol-rich membrane microdomains. As the brain ages, membrane cholesterol levels exhibit alterations, although it remains unclear how these changes impact protein-protein interactions and downstream signaling. Herein, we studied the structural bases for the interaction between GABA_B_R and the KCC2 transporter, including their protein expression and distribution, and we compared data between young and aged rat cerebella. Also, we analyzed lipid profiles for both groups, and we used molecular dynamics simulations on three plasma membrane systems with different cholesterol concentrations, to further explore the GABA_B_R-transporter interaction. Based on our results, we report that a significant decrease in GABA_B2_ subunit expression occurs in the aged rat cerebella. After performing a comparative co-immunoprecipitation analysis, we confirm that GABA_B_R and KCC2 form a protein complex in adult and aged rat cerebella, although their interaction levels are reduced substantially as the cerebellum ages. On the other hand, our lipid analyses reveal a significant increase in cholesterol and sphingomyelin levels of the aged cerebella. Finally, we used the Martini coarse-grained model to conduct molecular dynamics simulations, from which we observed that membrane cholesterol concentrations can dictate whether the GABA_B_R tail domains physically establish G protein-independent contacts with a transporter, and the timing when those associations eventually occur. Taken together, our findings illustrate how age-related alterations in membrane cholesterol levels affect protein-protein interactions, and how they could play a crucial role in regulating GABA_B_R’s interactome-mediated signaling.

**Significance Statement:** This study elucidates age-related changes in cerebellar GABA_B_ receptors (GABA_B_Rs), KCC2, and plasma membrane lipids, shedding light on mechanisms underlying neurological decline. Molecular dynamics simulations reveal how membrane lipids influence protein-protein interactions, offering insights into age-related neurodegeneration. The findings underscore the broader impact of cerebellar aging on motor functions, cognition, and emotional processing in the elderly. By elucidating plasma membrane regulation and GABAergic dynamics, this research lays the groundwork for understanding aging-related neurological disorders and inspires further investigation into therapeutic interventions.

## Introduction

Under normal aging processes, brain cells progressively decline and are lost, which leads to permanent deficits in motor, cognitive, and behavioral abilities (1). Although most of the underlying mechanisms remain elusive, almost all age-related neurodegenerative processes appear to be linked to any of the following alterations: 1) changes in membrane lipid composition, especially cholesterol, gangliosides, sphingolipids, and ceramides levels, which are the most affected by aging (2–8); 2) the inability to degrade cellular residues that accumulate in both inside and outside the cells (9–11); or 3) an imbalance between excitatory and inhibitory neural responses (12). These processes might not affect all brain areas at the same time or in similar ways. In the cerebellum, for example, variations in cholesterol and phospholipid compositions have been linked to aging (2, 4, 13, 14). Lipofuscin deposits have been found within aged Purkinje cell somata (15, 16). These accumulations, along with an unbalanced inhibitory signaling, account for some of the age-associated changes in the cerebellar tissue (17–21). Since the cerebellum is well integrated with other brain regions, age-induced changes that affect cerebellar circuits and its homeostasis could ultimately disturb motor functions, attention and learning, and the processing of language and emotions (22).

GABA_B_ receptors (GABA_B_Rs) are G-protein coupled receptors (GPCRs) that require the heterodimerization of GABA_B1_ and GABA_B2_ subunits, in order to establish functional complexes at the cell surface (23–25). Unlike GABA_A_ receptors (GABA_A_Rs), which are chloride-permeable ligand-gated ion channels that mediate fast responses to GABA (26, 27), metabotropic GABA_B_R actions are responsible for slower effects. Briefly, the binding of the inhibitory neurotransmitter GABA (gamma-aminobutyric acid) to GABA_B_Rs triggers activation of trimeric G_o/i_-type proteins, which in turn block cAMP production (28, 29). GABA also inhibits voltage-gated Ca^2+^ (Ca_v_) channels both presynaptically, preventing neurotransmitter release, and postsynaptically, hampering calcium-dependent dendritic spikes (30). In addition, it induces the opening of G protein-coupled inwardly rectifying K^+^ channels (GIRK) and the firing of slow inhibitory postsynaptic potentials (IPSP) (28, 29). These events lead to membrane hyperpolarization. For GABA_B1_, transcripts for several isoforms have been identified (31). In the cerebellum of adult rats, functional isoforms GABA_B1a-d_ along with GABA_B1f_, have been described (31–35). Once established at the cell surface, GABA_B_Rs associate with a broad range of proteins (28, 36, 37). GABA_B_Rs have been shown to interact with KCC2, an important neuronal symporter that actively extrudes chloride ions out of the cell by coupling their transport to the K^+^ electrochemical gradient (38, 39). There are two isoforms for the transporter, KCC2a and KCC2b, which differ mainly in their N-terminal region and form either monomers, homo- and heterodimers, or larger oligomers at the cell surface (38–42). KCC2 plays a fundamental role in the GABAergic system, since it is responsible for the maintenance of chloride gradients in mature neurons and, therefore, exerts a direct influence on the GABA_A_R driving force (43–45). GABA_B_Rs multiprotein complexes, as well as other cell-surface receptors like GABA_A_Rs, are often organized in lipid rafts enriched in cholesterol and sphingolipids (46–49). However, little is known about the influence of membrane lipid changes on protein dynamic organization at the cell surface and receptor-mediated signaling, especially regarding membrane alterations in aged tissue (8).

In the cerebellum, GABA_B_Rs and KCC2 are expressed widely in all layers of the cortex (50–52). In the molecular layer (ML), GABA_B1_ and GABA_B2_ subunits are distributed extensively through the Purkinje cell (PkC) dendritic trees (53). Higher densities are found at the periphery of excitatory synapses between parallel fibers (PFs) and PkCs (53–58). Regarding KCC2, it predominates in the distal and central regions of the ML. KCC2 is also expressed in adult cerebellar PkCs, where the transporter is found delineating the somatic membranes and the densely abundant PkC dendrites that are orientated towards PF synapses (51, 52, 59). In the ML, KCC2 is also expressed in GABAergic interneurons, specifically basket and stellate cells (60, 61). On the other hand, in the granular layer (GL) of the adult cerebellum, the expression of KCC2 is prominent, where it is present in both granule neurons and Golgi type II cells (59–62).

In this study, we aimed to investigate how healthy aging affects the integration of cerebellar GABAergic responses. We focused on interactions at the molecular level among GABA_B_Rs, KCC2, and membrane lipids. By applying analytical tools, including Western blot, ultra-high-performance liquid chromatography coupled to tandem mass spectrometry, and immunohistochemical analyses, we identified GABA_B_Rs and KCC2 subunits and isoforms, along with their expression levels and distribution patterns in the aged rat cerebellum. Among our findings, we discovered several GABA_B1_ and KCC2 isoforms, and we found that GABA_B2_ expression dropped progressively as the cerebellum aged. We also identified other important structural changes associated with aging, especially the accumulation of lipofuscin deposits inside and surrounding PkCs. Additionally, we used co-immunoprecipitation assays to confirm that GABA_B_Rs interact with KCC2 in the aged cerebella. Although, interaction levels were significantly lower than those detected in young adults. Moreover, we comparatively analyzed lipid composition profiles and found that cholesterol and sphingomyelin levels were substantially elevated in the aged cerebella. With these results in mind, we used coarse-grained molecular dynamics (CGMD) simulations to further investigate how membrane lipids impact the GABA_B_Rs dynamics and interactions. In three different lipid bilayer systems, where cholesterol concentrations covered 45%, 25%, and 10% of total lipid composition, we studied GABA_B_R’s conformational transitions, and GABA_B_R interactions with a KCC2-*like* transporter and the surrounding lipids. We discovered that changes in the global lipid configuration affected differently the timing and nature of protein-protein and protein-lipid interactions among GABA_B_Rs tail domains, the transporter, and lipids of the cytofacial layer. Additionally, in membrane systems with cholesterol concentrations of 45% and 10%, we identified a direct interaction between the GABA_B1_ subunit and the transporter. Overall, our results exposed numerous changes that occur during aging of the rat cerebellum, that impact lipid composition, while also affecting GABA_B_R and KCC2 expression and interactions.

## Results

### GABA_B_R subunits and KCC2 are found in multiple isoforms in the aged rat cerebellum

In this study, we aimed to better characterize protein expression and distribution for the metabotropic GABA_B_ receptor subunits along with KCC2, in the cerebellum of healthy aged rats. First, we solved total protein extracts from 18-month-old rat cerebella (P18m) and compared band profiles and protein levels to the ones acquired from young adult rats (P3m)

(Fig. 1A and 1B). Bands were detected for GABA_B1a_, GABA_B1b,_ GABA_B2_, and KCC2 (Fig. 1A). Quantification of band intensities revealed a significant drop for GABA_B2_ subunit levels from P3m to P18m, whereas GABA_B1_ isoforms did not present notable differences between ages (Fig. 1B). We observed substantial differences between dimeric and monomeric forms of KCC2, but their levels can vary depending on the buffer and protocol used for protein extraction (Fig. 1B). In fact, no significant differences were detected for total KCC2 expression between the young and aged groups. However, since all three assayed blots displayed multi-band profiles (Fig. 1A) and based on the evidence that several transcripts have been described for GABA_B1_ and KCC2, we performed UHPLC-MS/MS, followed by bioinformatics analysis, to identify the putative isoforms that are present in the aged cerebellum (P18m). The applied methodology is summarized in Figure 1C (Fig. 1C).

**Figure 1.**
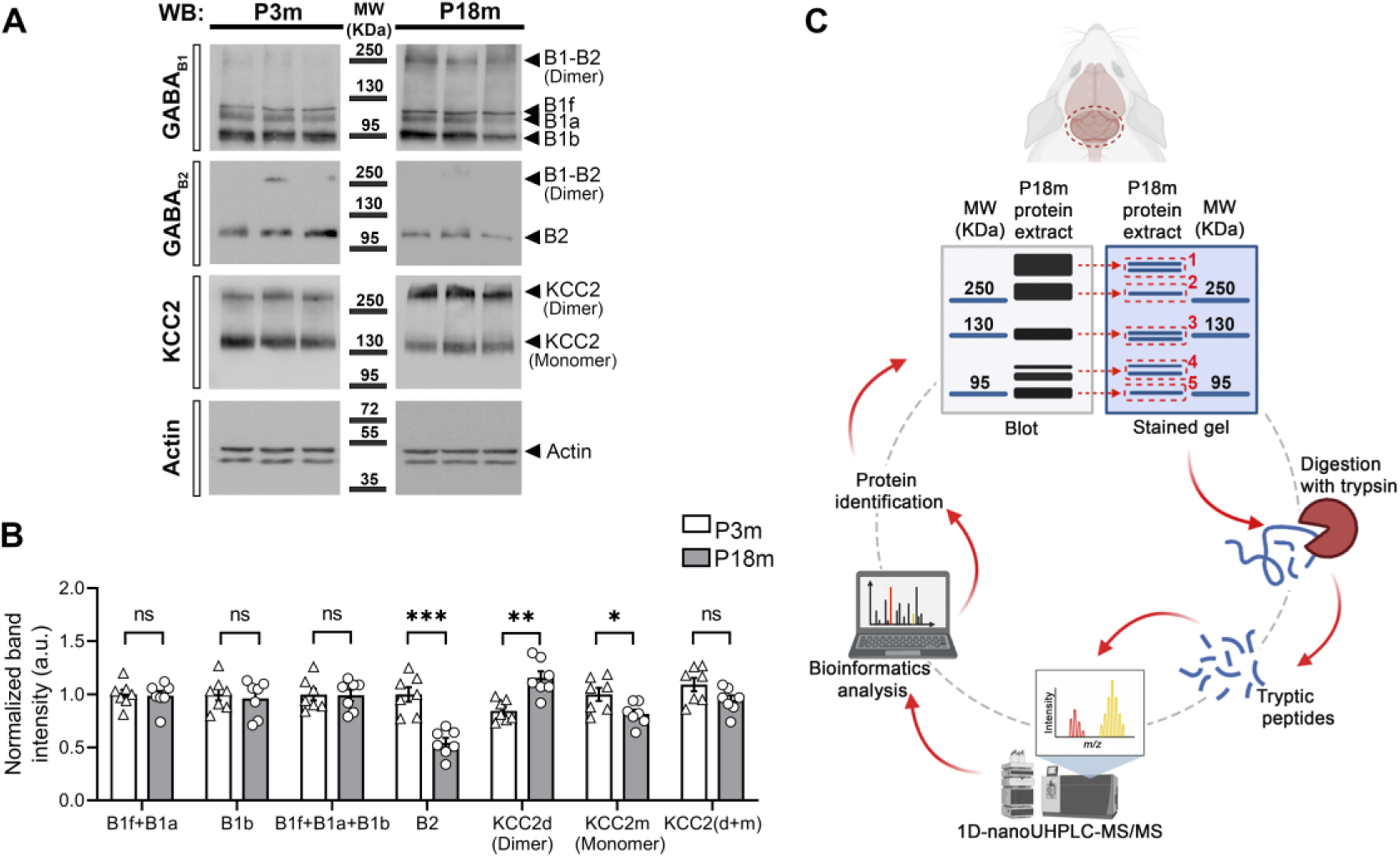
In P18m rat cerebellum, multiple isoforms are detected for GABA_B_R subunits and KCC2. Total protein expression comparison by Western blot (WB) between young adult and aged rat cerebella revealed multi-band profiles for the GABA_B_R subunits and KCC2. Also, GABA_B2_ expression declined significantly in the aged cerebella (\*\*\**p* = 0.0001). Total KCC2 expression [KCC2(d+m)] did not present significant variations although the dimeric form (d) was more abundant in the aged group (\*\**p* = 0.0012), while the monomeric form (m) prevailed in the young adult cerebella (\**p* = 0.0266). **A)** Representative blots display triplicates of protein extracts collected from P3m and P18m rat cerebella (3- and 18-month-old, respectively). Samples were generated using Triton X-100 lysis buffer. B1-B2 (Dimer): SDS-resistant GABA_B1_-GABA_B2_ heterodimer (∼280 kDa). B1f: GABA_B1f_ subunit (∼111 kDa). B1a: GABA_B1a_ (∼108 kDa). B1b: GABA_B1b_ (∼95 kDa). B2: GABA_B2_ (∼106 kDa). KCC2 (Dimer): ∼280 kDa. KCC2 (Monomer): monomeric forms of KCC2a and KCC2b (∼127 and ∼124 kDa, respectively) that weigh around 140 kDa when glycosylated. MW: molecular weight markers. **B)** Comparison of relative protein levels. Band intensities were normalized to total protein per lane. The graph exhibits mean ± SEM; n=7; a.u.: arbitrary units; ns: not significant. The Shapiro-Wilk normality test, followed by unpaired T-test, was applied for statistical analysis. KCC2(d+m): total KCC2 expression from dimeric and monomeric forms. **C)** The image summarizes the procedure applied to identify protein bands previously detected by WB. Briefly, total protein extracts from aged cerebella were generated using two different extraction buffers (NP-40 and Triton X-100 lysis buffers). Proteins were separated by gel electrophoresis and later stained with Coomassie Brilliant Blue G250. Then, five fragments were excised and trypsin-digested from gel regions where proteins of interest potentially migrated. Tryptic peptides were separated and identified by UHPLC-MS/MS, followed by bioinformatics analysis.

Using two distinct lysis buffers to generate protein extract facilitated the bioinformatic retrieval of different peptide pools from the same P18m cerebellar tissue. Also, as a first MS/MS analysis, each triplicate sample and its peptides were examined separately. Then, a second analysis was performed in which triads were unified and further treated as single samples. This comparative strategy allowed the proteomic information to be double confirmed and contrasted to determine any differences. Proteins identified in fragments 1-5, from both NP-40 and Triton X-100 lysis buffers, are listed in Table 1 and Dataset S1, along with the following descriptive data: UniProt accession number, a software-assigned score based on the Mascot server threshold for reliable protein identification, the number of retrieved peptides, the percentage of the given protein sequence covered by the retrieved peptides, protein isoelectric point (pl), an example of a peptide that unambiguously identifies the cited protein, and the supplementary dataset where all peptides can be found. In the case of the GABA_B1_ subunit, we were able to identify peptides that unambiguously corroborated the identity of some isoforms, and these sequences are presented in black color in Table 1. In some other cases, peptides consistent with more than one isoform sequence are presented in gray. When a given peptide was found only in the first series of individual analyses, in one out of the three analyzed fragment triplicates, then the peptide sequence is presented in blue. Unless stated otherwise, further descriptions and general conclusions mostly refer to the second analyses, performed after triplicate samples were unified. In all cases, valid protein identification was defined as protein coverages that were greater than 40%.

**Table 1.**
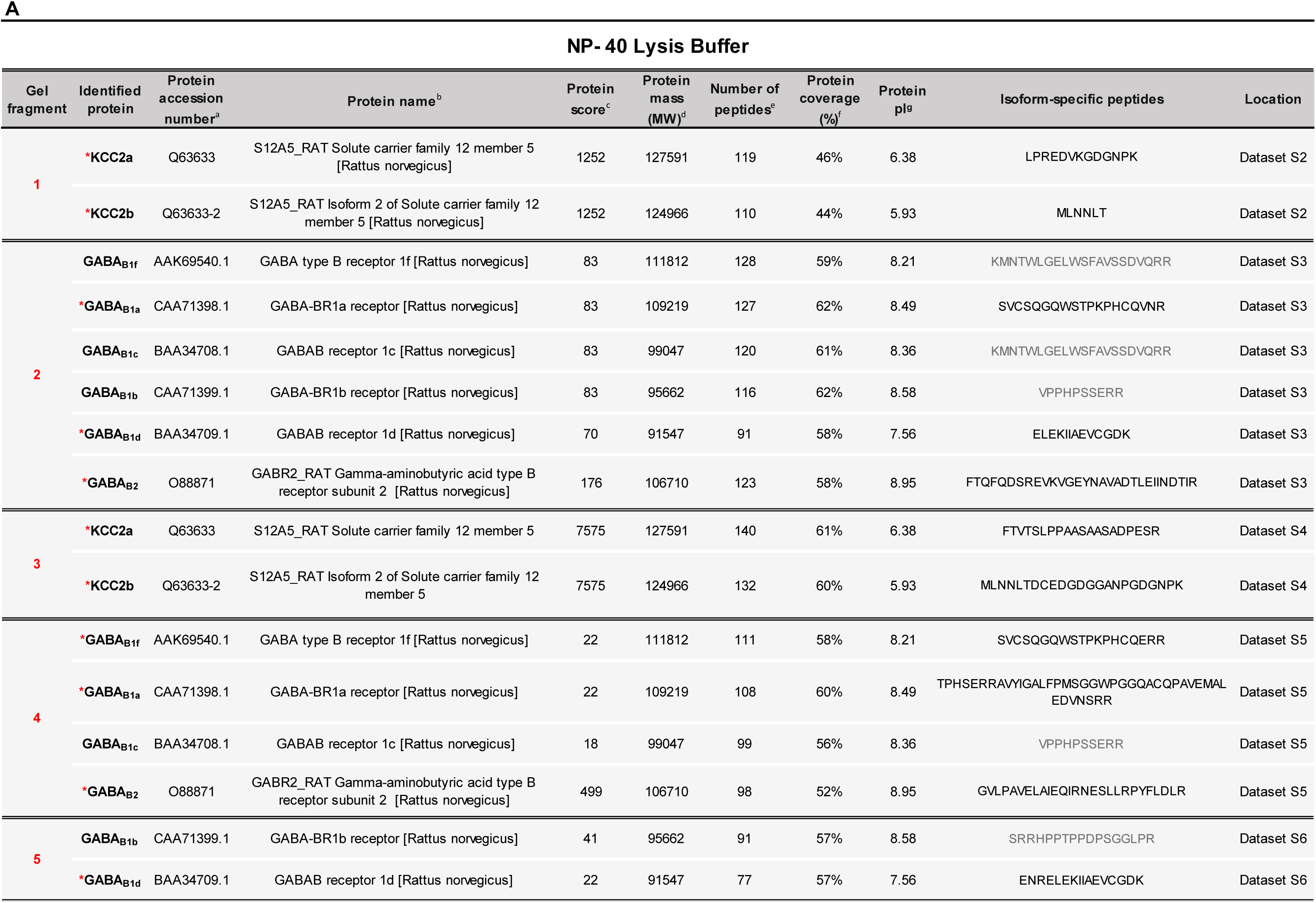
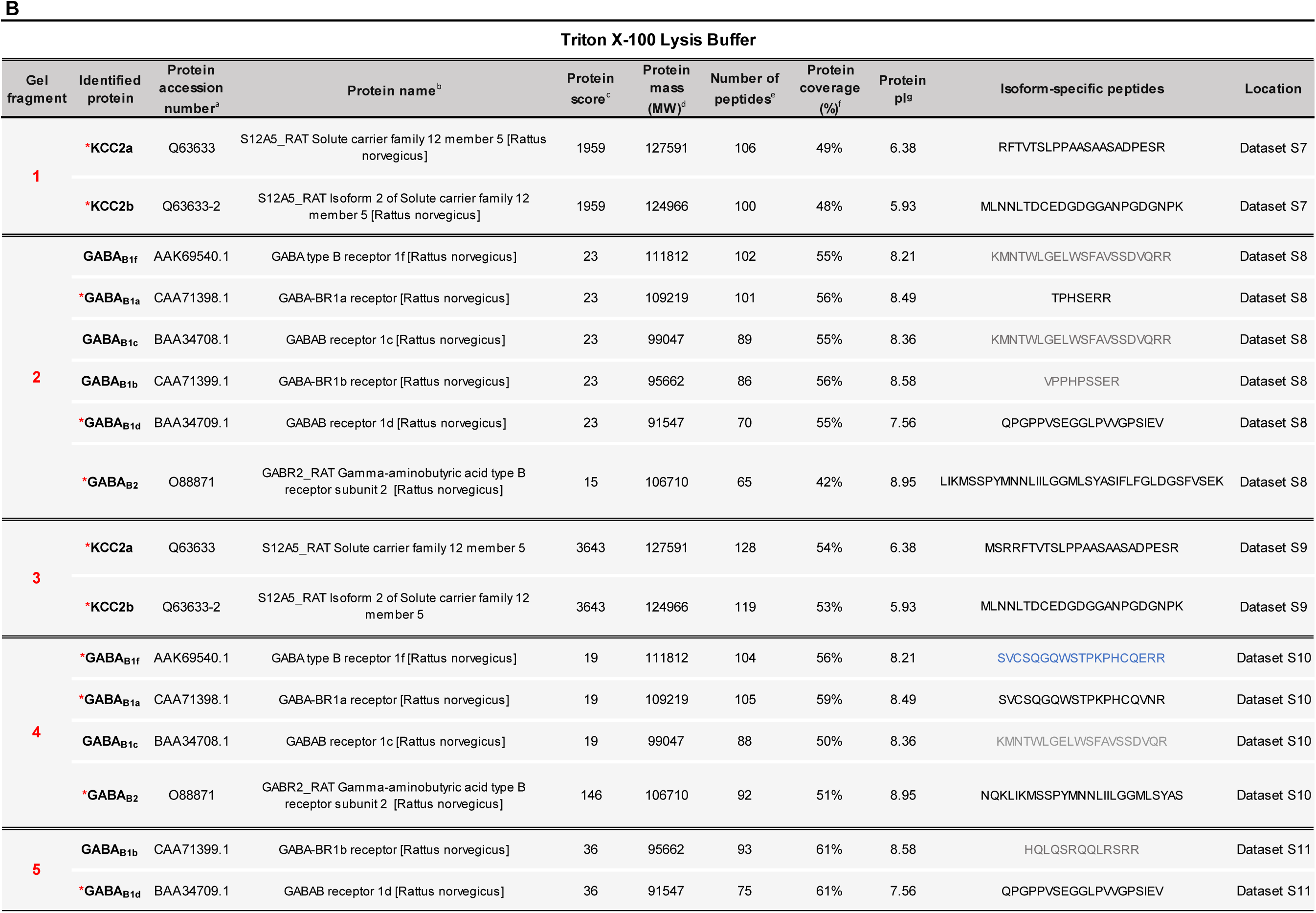
List of proteins and isoforms identified from each gel fragment by UHPLC-MS/MS analysis. Total protein extracts were generated using two lysis buffers: NP-40 **(A)** or Triton X-100 **(B).** Samples in triplicates were resolved by gel electrophoresis and treated as described in Fig. 1C. For each excised gel fragment, the identified proteins are listed, along with the following descriptive data: accession number, name, score, mass, number of retrieved peptides, protein coverage, protein isoelectric point, an example of an identified isoform-specific peptide, and the supplementary dataset to refer to. **(*)** Proteins unmistakably identified, for which isoform-specific peptides were detected. Peptide sequences that matched more than one GABA_B1_ isoform are exhibited in grey. In blue, the retrieved peptide paired to the GABA_B1f_ isoform was found only in one of the triplicates. **(a)** and **(b)** were retrieved from Swiss Protein (www.uniprot.org) and RefSeq (www.ncbi.nlm.nih.gov), respectively. **(c)** The Mascot server established the threshold setups for random hits contemplating a significance level at *p* ≤ 0.05. Those scores of greater thresholds were considered significant matches (www.matrixscience.com). **(d)** and **(g)** are the theoretical molecular weight (MW) and isoelectric point (pl), obtained from Swiss Protein (www.uniprot.org) and RefSeq (www.ncbi.nlm.nih.gov), respectively. **(e)** The number of matching peptides to the target protein, as corroborated by MS/MS analysis. **(f)** The percentage of a given protein sequence covered by all amino acids identified by MS/MS analysis from peptide mapping that successfully matched the designated protein.

From fragment 1, multiple peptides were recovered, some of which were consistent with a conserved and shared region between both KCC2 isoforms. In addition, we successfully identified peptides which sequences matched and differentially confirmed the presence of KCC2a (∼127 kDa) and KCC2b (∼124 kDa) within the analyzed samples (red asterisks; Table 1A1 and 1B1; Datasets S2 and S7). Based on the molecular weights of the transporter isoforms, and the identity of the recovered peptides, and considering the ability of KCC2 to organize into homo- or heterodimers, some of which are SDS-resistant (63), we named the high molecular weight band observed in KCC2 blots as KCC2 (Dimer) (Fig. 1A). Predictably, from fragment 3 peptide analyses, we also recovered specific peptides that allowed the recognition of the two KCC2 isoforms in the aged cerebellum (Table 1A3 and 1B3; Datasets S4 and S9).

For fragment 2, recovered peptides from each lysis buffer matched the sequences of both GABA_B1_ and GABA_B2_ subunits (Table 1A2 and 1B2; Datasets S3 and S8). For GABA_B1_, we identified isoform-specific peptides for GABA_B1a_ and GABA_B1d_ (red asterisks; Table 1A2 and 1B2). Moreover, peptides that paired shared regions between GABA_B1f_ and GABA_B1c_ (KMNTWLGELWSFAVSSDVQRR), or among GABA_B1b,_ GABA_B1c_, and GABA_B1d_ (VPPHPSSERR) were detected (in gray, Table 1A2 and 1B2). Considering the molecular weights of GABA_B_R subunits and isoforms (∼91-111kDa), peptides recovered at ∼250 kDa probably belonged to a fraction of stable GABA_B_R heterodimers which outlasted the denaturalization steps preceding protein electrophoresis (64–66). Therefore, we identified the higher bands observed in GABA_B1_ and GABA_B2_ blots as B1-B2 (Dimer) (Fig. 1A). Continuing with the analysis of GABA_B_R bands, fragment 4 provided peptides that specifically matched GABA_B2_, GABA_B1f_, and GABA_B1a_ sequences (red asterisks; Table 1A4 and 1B4; Datasets S5 and S10), along with multiple peptides consistent with regions shared among more than one GABA_B1_ isoform. Finally, from gel fragment 5, software analysis allowed the identification of GABA_B1b_ in the samples, but other peptides were also obtained that unmistakably confirmed the presence of GABA_B1d_ in the aged rat cerebellum (Table 1A5 and 1B5; Datasets S6 and S11).

Considering that GABA_B1a_ (∼108 kDa) and GABA_B1b_ (∼95 kDa) are the dominant GABA_B1_ isoforms in the brain (24, 32), we named the two lower bands observed in the GABA_B1_ blots as B1a and B1b, respectively (Fig. 1A). Furthermore, the thin band observed right above GABA_B1a_ (B1a) in the GABA_B1_ blots was designated as B1f, since we identified peptides matching the GABA_B1f_ isoform sequence in fragment 4 (Table 1A4 and 1B4). In addition, B1f is the only isoform with a theoretical weight (∼111 kDa) higher than B1a. Interestingly, in the KCC2-enriched gel fragments 1 and 3, we also retrieved peptides that matched the sequences of the GABA_B_R subunits and isoforms (Datasets S2, S4, S7 and S9). Consistently, for gel fragments 2, 4, and 5, where only GABA_B_R-related peptides were expected, we obtained several sequences that aligned with both KCC2 isoforms (Datasets S3, S5, S6, S8, S10 and S11).

### Protein levels of the GABA_B2_ subunit gradually decrease as the cerebellum ages

Since GABA_B2_ protein levels decayed in whole extracts from young (P3m) to older (P18m) cerebella (Fig.1A and 1B), we further explored changes in the subunit expression and distribution by applying multiple fluorescent immunolabeling followed by confocal microscopy. In this analysis we also included cerebella from 21-month-old rats (P21m), to better understand any age-related changes in the GABAergic system (Figs. 2 and S1). Additionally, we focused our study on the Purkinje cells (PkC) and their projections that are orientated towards the molecular layer (ML).

**Figure 2.**
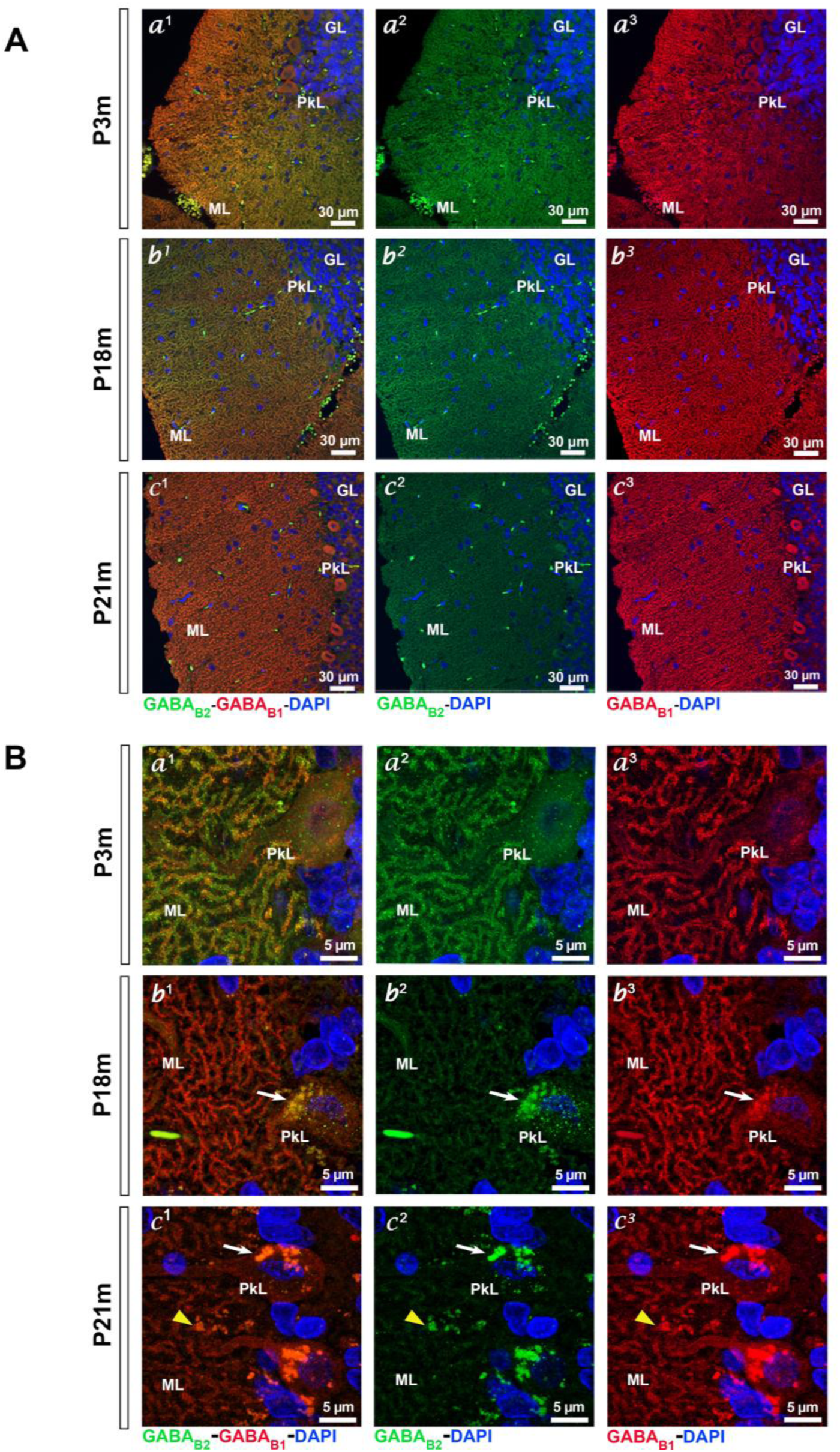
As the rat cerebellum ages, protein levels for the GABA_B2_ subunit progressively decrease in the molecular layer. Cerebellar sections from P3m (a^1^-a^3^), P18m (b^1^-b^3^) and P21m (c^1^-c^3^) rats were immunolabeled for GABA_B1_ (red) and GABA_B2_ (green). DAPI (blue) was used as a nuclear marker. **A)** Images at 60X show GABA_B1_ and GABA_B2_ expression and distribution in the Purkinje cell and molecular layers (PkL and ML, respectively). GABA_B2_ signal gradually dimed from a^2^ to b ^2^, and it became barely detectable at c^2^, which confirmed a decreasing gradient as age progressed. GL: granular layer. **B)** 2.1x digital zooms from 100X images evidenced an overlapping pattern for GABA_B_R subunits in the PkL and ML proximal region of the P3m cerebellum (a^1^). With age, GABA_B2_ fluorescence faded out until the Purkinje cell (PkC) dendritic tree was no longer recognizable (c^2^). At P18m, fluorescent lipofuscin-like small granules emerged within the PkC cytoplasm (white arrow, b^1^-b^3^). At P21m, larger accumulations were detected inside the PkC somata (white arrow, c^1^-c^3^) and in the PkC periphery (yellow arrowhead, c^1^-c^3^).

In the young adult cerebellum (P3m), we detected that GABA_B1_ and GABA_B2_ were highly expressed on the PkC somata and throughout the ramifications of their dendritic trees, where both subunits were often found to be colocalized (Fig. 2A a^1^). However, colocalization diminished as the cerebellum aged (Fig. 2A b^1^ and c^1^), which was caused by a global and gradual decrease in the GABA_B2_ signal. A significant reduction of the GABA_B2_ immunoreactivity was observed in P18m cerebellar sections, and this fade-out effect was noticeably exacerbated in P21m cerebella (Fig. 2A b^2^ and c^2^). This led us to conclude that the phenomenon is progressive and seems to be associated with the aging process.

In a deeper analysis of the Purkinje cell layer (PkL) and their nearby areas, we observed substantial signal overlapping of GABA_B1_ and GABA_B2_ in dendritic tree membranes of P3m cerebella (Fig. 2B a^1^-a^3^), as has also been described elsewhere (67, 68). In the older P18m and P21m stages, the granular pattern first observed in the young P3m ML, progressively disappeared as the GABA_B2_ immunoreactivity declined (Fig. 2B b^2^ and c^2^). On the other hand, GABA_B1_ presented a robust signal on PkC somata and along the intricated branches of their dendritic trees, and the observed intensity and distribution did not significantly fluctuate throughout the adulthood stages analyzed in this study (Fig. 2B a^3^, b^3^ and c^3^).

Additionally, we detected other morphological changes emerging in the older rat cerebellum. Discrete fluorescent aggregates were observed in P18m sections inside the PkC somas (white arrow; Fig. 2B b^1^-b^3^). Similarly, we identified irregular assemblies in P21m, although they were located both within and around PkCs (white arrow and yellow arrowhead, respectively; Fig. 2B c^1^-c^3^). Intracellular accumulations have previously been described in the aged cerebellum as deposits of lipofuscin (LP), and they are generally considered to be a hallmark of aging (9, 15, 69). Moreover, autofluorescence is one of the main LP characteristics (9, 70). Hence, we scanned cerebellar sections across different wavelengths to detect fluorescent emission. By applying this methodology, we were able to recognize autofluorescent LP aggregates both inside and outside of P21m PkC somata (white arrows and yellow arrowhead, respectively; Fig. S1 a^1^-d^3^). At this P21m stage, we also detected bright deposits in the cytoplasm of other cells of the ML (white arrowhead; Fig. S1 b^1^-b^3^). In conclusion, the presence of lipofuscin clusters confirmed the impact of healthy aging upon the cerebellar architecture and its cellular integrity.

### KCC2 is expressed in the PkC membrane, but the distribution patterns change as the cerebellum ages

To better characterize GABA_B_R location related to KCC2 in the cerebellum, we immunolabeled P3m and P18m cerebellar sections for GABA_B1_ and KCC2 proteins. Then, we investigated the spatial distribution patterns and potential overlapping of both molecules in the ML and PkL (Fig. 3A and 3B). This analysis revealed an extensive signal for KCC2 in both stages, although with subtle differences. In the PkL and proximal ML of the young cerebellum, KCC2 was detected in the perimeter of PkC somata and throughout the membranes of their projections (white arrowheads; Fig. 3A a^1^ and a^2^). In the distal P3m ML, KCC2 staining intensively overlapped the GABA_B1_ signal along membrane processes (white arrowhead; Fig. 3B a^1^-a^3^). However, high KCC2 immunoreactivity was also observed where GABA_B1_ was undetectable, as in widely distributed thinner branches and smaller-sized cells (yellow arrowheads; Fig. 3A a^1^-a^3^ and 3B a^1^-a^3^). Overall, KCC2 cellular and subcellular distribution was substantially the same in both young and aged cerebella, except for minor variations. Even though no statistically significant difference was detected in global KCC2(d+m) levels between P3m and P18m via WB (Fig. 1A and 1B), we observed a slight attenuation for KCC2 immunoreactivity around PkC somata and their dendritic trees in the P18m cerebellum (white arrowhead; Fig. 3A b^1^ and b^2^). However, high-intensity KCC2 signal observed in small interneurons seemed to be unaltered in the P18m ML (yellow arrowhead; Fig. 3A b^1^ and b^2^). In the P18m PkL, we confirmed age-related changes by the detected presence of intracytoplasmic LP deposits (white arrow; Fig. 3A b^1^-b^3^). Similarly, in the distal P18m ML, we observed a reduction of KCC2 staining in GABA_B1_-positive branches which was accompanied with a more clumped membrane distribution of the transporter (Fig. 3B b^1^-b^3^). Conversely, the intense KCC2 immunoreactivity observed in narrow membrane-delimited projections appeared to be unaffected, when distal MLs from young and aged cerebella were compared (yellow arrowheads; Fig. 3B a^1^- a^2^ and b^1^-b^2^).

**Figure 3.**
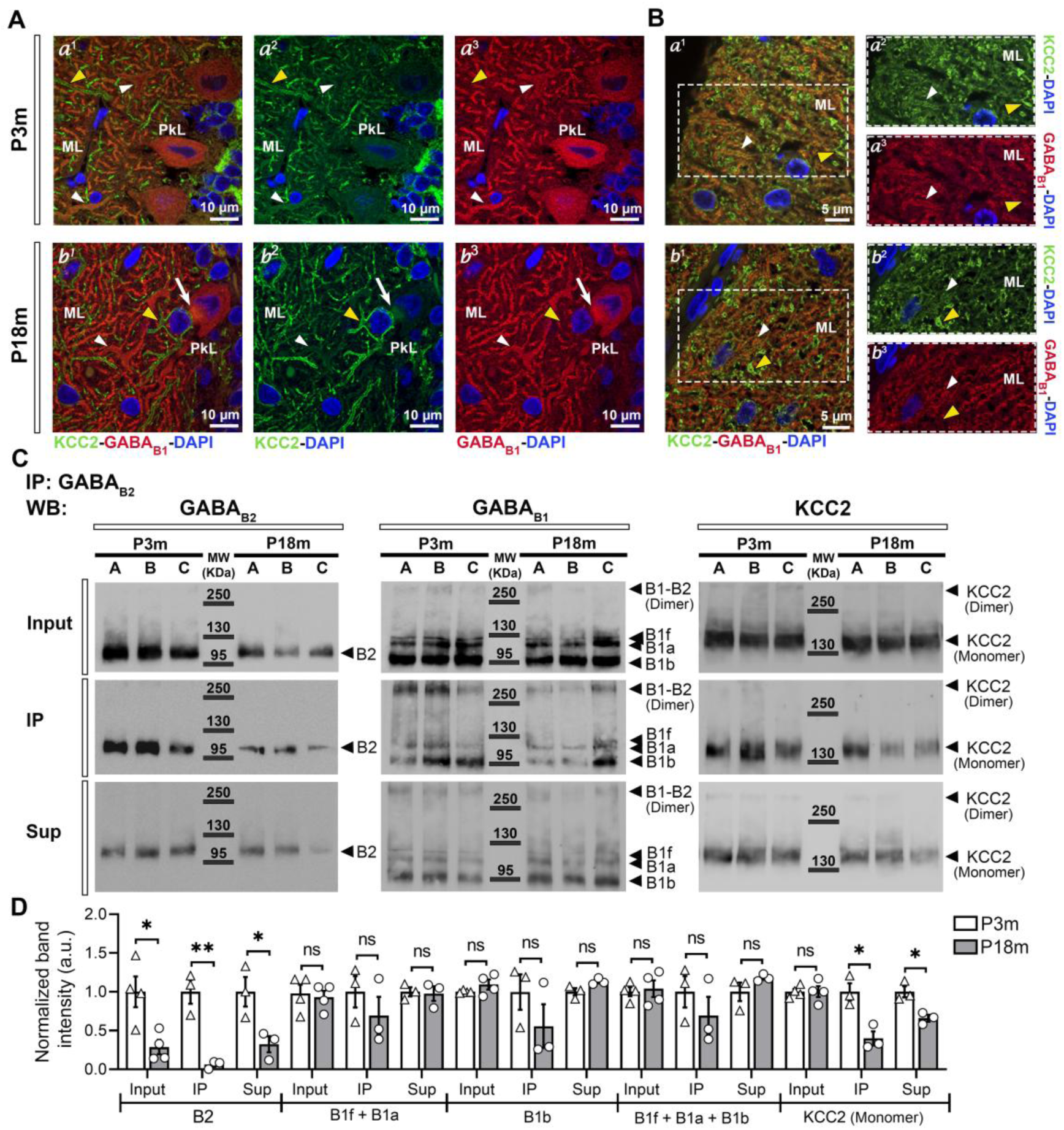
The GABA_B_ receptor and KCC2 interact and form a multiprotein complex in young adult and old rat cerebella, but the magnitude of this protein-protein interaction significantly decreases with aging. **A-B)** Confocal microscopy was applied to study cerebellar sections from P3m (a^1^-a^3^) and P18m (b^1^-b^3^) rats. Images exhibit immunoreactivity for KCC2 (green) and GABA_B1_ (red) in the Purkinje cell and molecular layers (PkL and ML, respectively). DAPI (blue) was used as a nuclear marker. **A)** Intense immunoreactivity for GABA_B1_ is detected in the Purkinje cell (PkC) somata and along their dendritic trees in young and aged cerebella (a^3^ and b^3^). KCC2 is expressed in the PkC dendrites and in the dendritic trunks as well. However, compared to the GABA_B1_ signal, a more subtle mark is observed for the transporter, which appears to be restricted to the plasma membrane (white arrowheads, a^1^-a^2^ and b^1^-b^2^). A small-sized cell population and multiple projections present immunoreactivity only for KCC2 in the proximal ML (yellow arrowheads, a^1^-a^2^ and b^1^-b^2^). Fluorescent lipofuscin aggregates appear in the soma of PkCs at 18 months (white arrow). 100X images with a 1.5x digital zoom. **B)** In the distal ML, an overlapping distribution pattern throughout the PkC dendritic membranes is observed for GABA_B1_ and KCC2; a reduction in signal colocalization and a transporter redistribution are detected in the aged cerebellum (white arrowheads, a^1^-a^3^ and b^1^-b^3^). Cells and projections immunoreactive only for KCC2, are also detected in this ML region (yellow arrowheads, a^1^-a^2^ and b^1^-b^2^). 100X images with a 2.0x digital zoom. **C)** Co-immunoprecipitation, followed by WB, was performed in protein extracts from P3m and P18m rat cerebella, by using the NP-40 lysis buffer and an anti-GABA_B2_ antibody to preserve protein-protein interactions and to isolate multiprotein complexes. After the pull-down assay, a fraction of KCC2 was retrieved from the immunoprecipitation product (IP), along with GABA_B2_ and GABA_B1_. Input: total protein lysate. Sup: remaining supernatant. B1-B2 (Dimer): SDS-resistant GABA_B1_ and GABA_B2_ heterodimer (∼280 kDa). B1f: GABA_B1f_ subunit (∼111 kDa). B1a: GABA_B1a_ (∼108 kDa). B1b: GABA_B1b_ (∼95 kDa). B2: GABA_B2_ (∼106 kDa). KCC2 (Dimer): ∼280 kDa. KCC2 (Monomer): monomeric forms of KCC2a and KCC2b (∼127 and ∼124 kDa, respectively) that weigh around 140 kDa when glycosylated. MW: molecular weight markers. **D)** Quantitative analysis of band intensities, which were normalized to total protein per lane. The graph exhibits mean ± SEM; n=3 (A-C); a.u.: arbitrary units; ns: not significant. The Shapiro-Wilk normality test, followed by unpaired T-test, was applied for statistical analysis. The experiment confirmed that GABA_B2_ expression diminishes as the cerebellum ages (Input: \**p* = 0.016703; IP: \*\**p* = 0.003922; Sup: \**p* = 0.035535). Levels of monomeric KCC2 in IP and Sup fractions were significantly lower in P18m, as compared to P3m samples (IP: \**p* = 0.013536; Sup: \**p* = 0.015058). A decreasing tendency was observed for the GABA_B1_ subunits when P3m and P18m IPs were compared.

### GABA_B_R and KCC2 interact in the cerebellum, but the formation of this protein complex is affected by aging

Previous published studies have reported GABA_B_R-KCC2 interactions in the rat cortex and hippocampus (39), as well as in mouse forebrain-derived neuronal plasma membranes, and cortical and hippocampal cell cultures (38). As described above, we also observed partial overlapping of GABA_B1_ and KCC2 on the PkC somata and their dendritic trees in the cerebellum (Fig. 3A and 3B). Based on these findings, we investigated further to determine whether a similar GABA_B_R-KCC2 association takes place in the cerebellum. Hence, we performed co-immunoprecipitation assays in total protein extracts from P3m and P18m cerebella. We used antibodies against GABA_B1_, GABA_B2_, and KCC2, to isolate immunocomplexes. Our immunoblotting analyses of each immunoprecipitation product (IP) confirmed that a fraction of GABA_B_R subunits along with KCC2 established a multiprotein complex in the cerebellum at both ages (Figs. 3C, S2 and S3). Considering the gradual decay of the GABA_B2_ subunit that occurs as the cerebellum ages (Fig. 2), we investigated whether aging also affected the GABA_B_R-KCC2 association. For this purpose, we performed a semiquantitative co-immunoprecipitation assay using an anti-GABA_B2_ antibody to compare levels of proteins isolated from P3m and P18m multiprotein complexes (Figs. 3C, 3D and S2). We detected significantly lower GABA_B2_ in the input, the pull-down product, and the remaining supernatant from P18m cerebellar extracts, as compared to GABA_B2_ levels in the P3m fractions (Fig. 3D). In addition, we found that GABA_B1_ and KCC2 co-immunoprecipitated with GABA_B2_ when protein complexes were isolated from both stages. GABA_B1_ levels detected in the different fractions did not present statistically significant differences between P3m and P18m. However, we observed a decreasing tendency for GABA_B1_ subunits (GABA_B1f_ + GABA_B1a_, GABA_B1b_, and GABA_B1f_ + GABA_B1a_ + GABA_B1b_) in the P18m immunoprecipitation product, as compared to P3m IP (Fig. 3D). On the other hand, levels of monomeric KCC2 retrieved from P18m IP and Sup fractions were significantly reduced, as compared to the ones obtained from P3m samples (Fig. 3D). Interestingly, KCC2 was detected mainly as a monomer when the anti-GABA_B2_ and anti-GABA_B1_ antibodies were used to isolate protein complexes (Figs. 3C and S2A).

In the cerebellum, KCC2 is in pre- and post-synaptic membranes at inhibitory neuronal junctions, where it may interact with the ionotropic GABA_A_Rs (38, 71). Therefore, we investigated the potential role of KCC2 as a linker between GABA_B_Rs and GABA_A_Rs in multiprotein complexes. We selected the α1 subunit since it is widely distributed in the cerebellum, where is part of the pentameric GABA_A_Rs found in granule and Purkinje cells (72–74). Consequently, we searched for the GABA_Aα1_ subunit in the pull-down fractions obtained from immunoprecipitation reactions performed with anti-GABA_B2_ and anti-GABA_B1_ antibodies. Our results confirmed that small fractions of α1 were present in the immunoblots from P3m and P18m IPs (Fig. S4). Pull-down fractions generated with the anti-KCC2 antibody were excluded from this analysis due to potential interference in blotting detection and antibody cross-reactions. This strategy took into consideration that the molecular weight of the α1 subunit matches the size of the heavy chain of an antibody (∼50 kDa), and both the highly specific anti-KCC2 and anti-GABA_Aα1_ antibodies used in the co-immunoprecipitation assay were generated in the same species.

### Cholesterol concentrations and sphingomyelin abundance are altered in the aged rat cerebellum

Changes in the composition of membrane lipids, including cholesterol, have been associated with aging and aging-related neurodegenerative processes. However, up to date, brain lipid profiles and variations seem to rely on the analyzed models, cerebral regions, cell types, sex, and age, among other variables (2, 4, 6, 13, 14, 75–80). Based on our results of age-associated changes in protein profiles and protein-protein interactions for the GABAergic system (Figs.1A, 1B, 3C and 3D), we investigated whether aging could also affect the lipid composition of the rat cerebellum. We collected cerebella from P3m and P18m rats and performed total lipid extraction, followed by lipid analyses. Total cholesterol and free cholesterol levels were found to be significantly increased in the aged group, which was accompanied by a substantial reduction of esterified cholesterol (Fig. 4A, 4B, and 4C). On the other hand, the concentration of lipid phosphorus and, therefore, that of phospholipids, did not vary significantly with age (Fig. 4D and 4E). Notably, this resulted in an increased cholesterol/phospholipid ratio in the P18m cerebellar tissue (Fig. 4F). In addition, levels of individual phospholipid classes did not change extensively between the P3m and P18m groups, with the exception that sphingomyelin abundance increased moderately, but significantly, in the older cerebella (Fig. S5).

**Figure 4.**
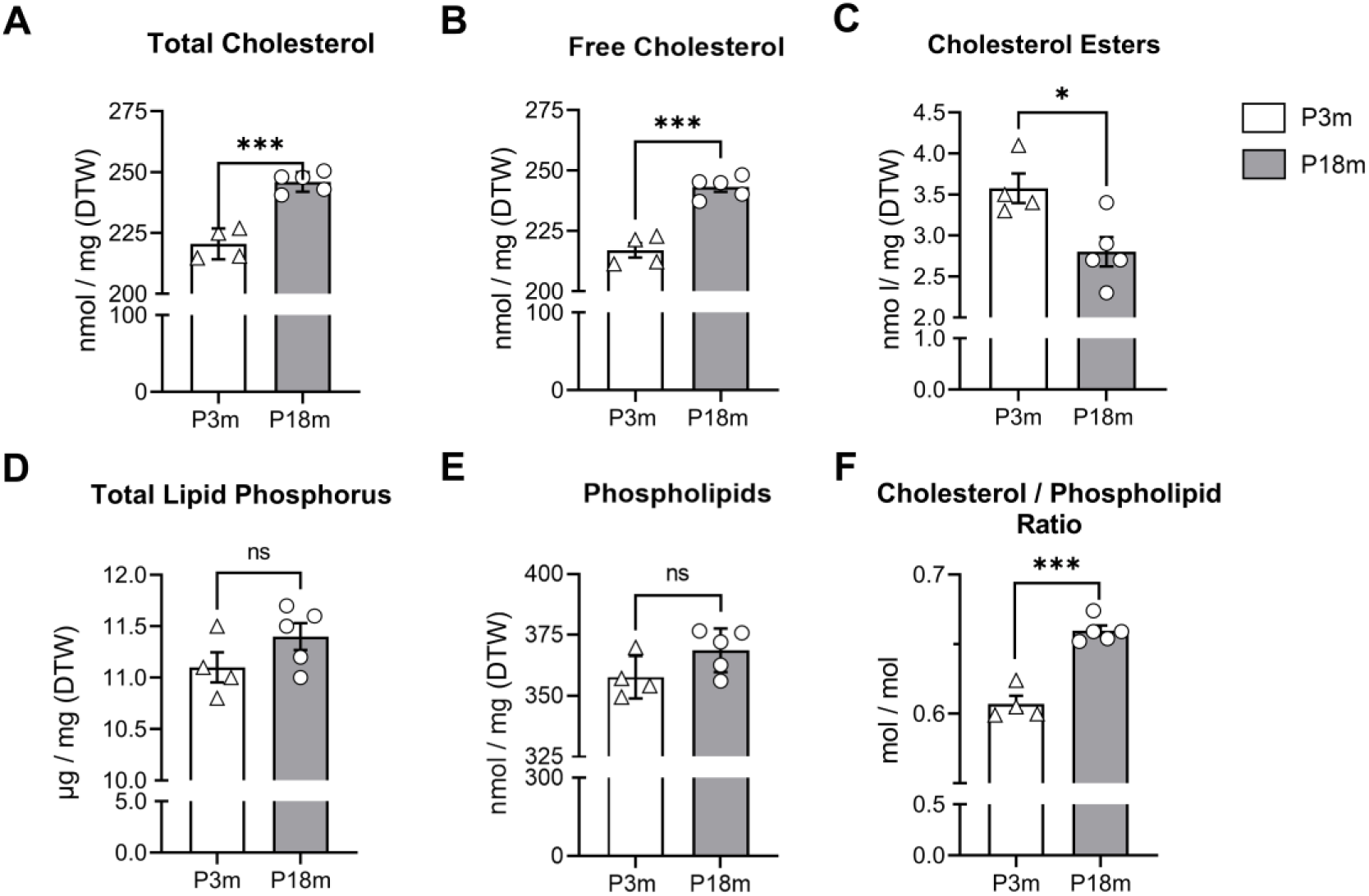
In the rat cerebellum, cholesterol levels increase significantly with age. Total lipid extracts from P3m and P18m rat cerebella were resolved by thin-layer chromatography (TLC). Comparison of relative lipid contents exhibited variable cholesterol concentrations between the P3m and P18m groups. Incremented amounts of total and free cholesterol were detected in the P18m group, as compared to levels measured in young adult rat cerebella. Likewise, the cholesterol/phospholipid ratio was significantly higher in the P18m samples. In contrast to all parameters mentioned before, cholesterol ester concentrations were substantially reduced in the older cerebella, as compared to P3m samples. Graphs exhibit means ± SEM; n=4 (P3m) and n=5 (P18m); ns: not significant. The Shapiro-Wilk normality test, followed by unpaired T-test, was applied for statistical analysis. **A)** Total cholesterol (\*\*\**p* = 0.0001). **B)** Free cholesterol (\*\*\**p* = 0.0001). **C)** Cholesterol esters (\**p* = 0.0125). **D)** Total lipid phosphorus (ns). **E)** Phospholipids (ns). **F)** Ratio of cholesterol content to phospholipids (\*\*\**p* = 0.0001). DTW: dry tissue weight.

### Coarse-grained molecular dynamics (CGMD) simulations allowed a deeper analysis of the GABA_B_R-KCC2 interaction

Studying GPCRs in their native biological membrane environment is challenging due to their structural and mechanistic complexity (81, 82). Molecular dynamics (MD) simulations have arisen in cell biology to complement experimental procedures and to provide fundamental insights into the dynamic properties and underlying mechanisms of interactive membrane proteins and lipids (81, 83, 84). Given the evidence that aging alters lipid profiles (especially cholesterol; Fig. 4) while also modifying protein levels and protein-protein interactions for the GABAergic system in the rat cerebellum (Figs. 1 and 3), we applied the Martini coarse-grained (CG) model to characterize more fully the interplay of GABA_B_Rs with other transmembrane proteins and surrounding lipids (Fig. 5).

**Figure 5.**
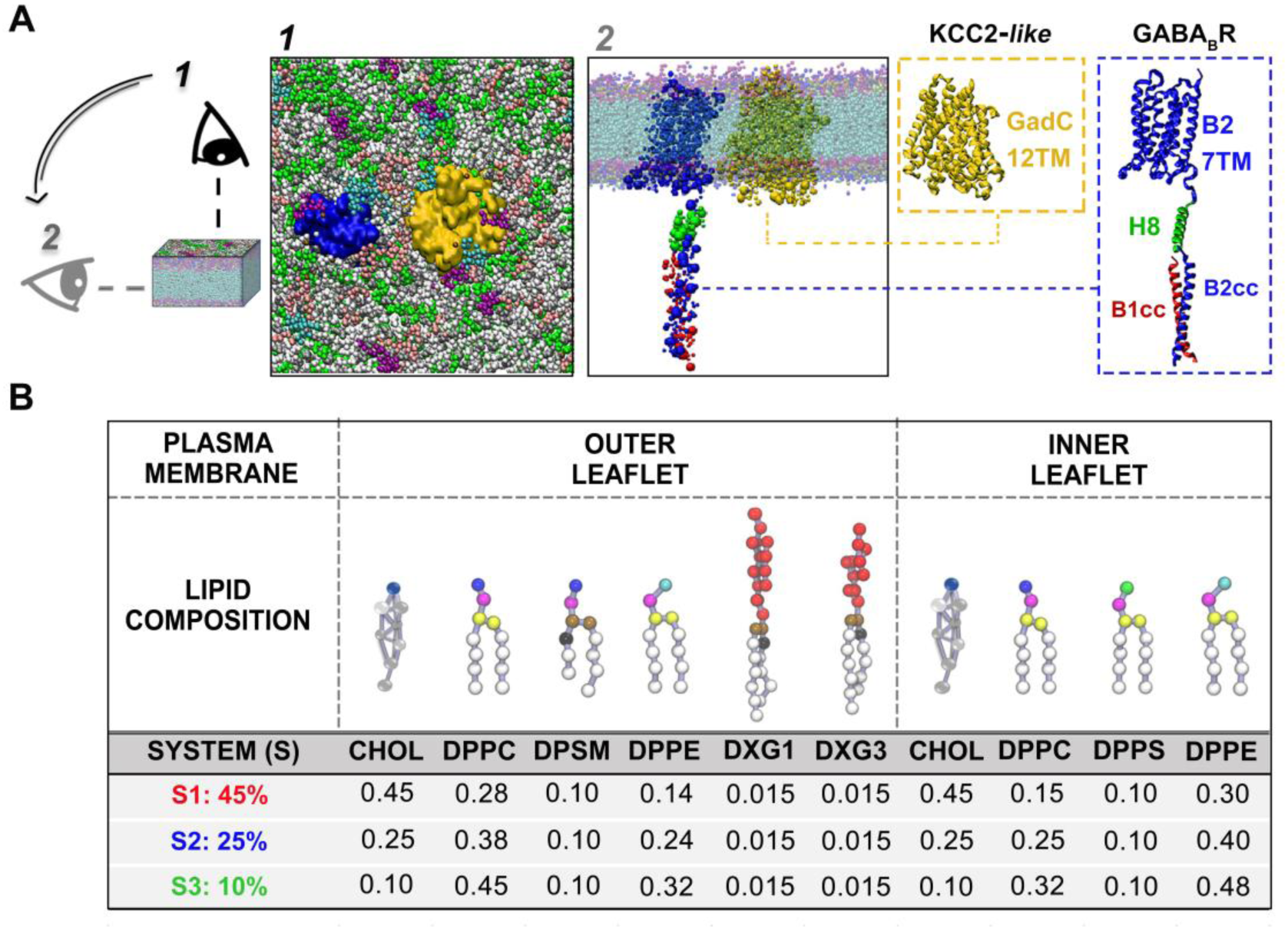
Coarse-grained molecular dynamics (CGMD) simulations of partial protein structures immersed in three plasma membrane system models were conducted to further study the GABA_B_R-KCC2 interaction. **A)** Martini CG model representations for a partial GABA_B_R (blue) and a KCC2*-like* protein (yellow), embedded in a lipid bilayer, are observed from two different points: 1) top view and 2) side view. In the blue inset, an atomistic model for the partial structure of GABA_B_R shows the domains selected for the simulations: the seven-transmembrane domain (7TM; blue), the alpha helix 8 (H8; green), and the coiled-coil domain from the human GABA_B2_ subunit (B2cc; blue), and the coiled-coil domain of the human GABA_B1_ subunit (B1cc; red). In the yellow inset, an atomistic representation is exhibited for the 12 alpha helixes of the bacterial glutamate-GABA antiporter (GadC; PDB ID: 4DJK). This structure was used for the simulations since it is homologous with the 12TM domain architecture of KCC2 and is referred to herein as KCC2*-like*. **B)** Three plasma membrane system (S) models with different lipid compositions, were simulated. For S1, S2, and S3, symmetrical cholesterol concentrations [CHOL] were 45%, 25%, and 10%, respectively, for both the inner and outer leaflets. DPPC and DPPE concentrations were modified proportionally to account for [CHOL] modifications. Martini topology models are presented for DPPC (dipalmitoylphosphatidylcholine: di-C16:0-C18:0 PC); DPPE (dipalmitoylphosphatidylethanolamine: di-C16:0-C18:0 PE); DPPS (dipalmitoylphosphatidylserine: di-C16:0-C18:0 PS); DPSM [sphingomyelin: C(d18:1/18:0) SM]; DXG1 [monosialotetrahexosylganglioside: C(d24:1/24:0) GM1]; DXG3 [monosialodihexosylganglioside: C(d24:1/24:0) GM3]. Models were retrieved from the Martini’s web page: www.cgmartini.nl/index.php/force-field-parameters/lipids.

For the construction of a computational plasma membrane (PM), we started with a general set of parameters for a typical human neuronal PM, as proposed by Ingólfsson *et al.* (2017) (85). We defined abundance of the major membrane lipid classes for both outer and inner leaflets, as the authors suggested. We designated this setup as System 1: 45% (S1: 45%), referring to the cholesterol percentage used (Fig. 5B). Considering that cholesterol was one of the most variable lipids in the rat cerebellum according to age (Fig. 4), we designed two other lipid bilayer systems where cholesterol concentrations [CHOL] were modified to represent 25% and 10% of the total lipids in each leaflet (S2: 25% and S3: 10%, respectively; Fig. 5B). The phospholipids DPPC and DPPE were adjusted proportionally to account for the designated variations in [CHOL]. Functional neuronal PMs exhibit an asymmetrical lipid distribution between the two leaflets (3, 86). In general, higher cholesterol concentrations are found at the cytofacial leaflet of the bilayer, whereas gangliosides are located at the exofacial layer of the neuronal PM (86). For our *in silico* assays, the systems were simplified with symmetrical [CHOL], while maintaining differential concentrations and distributions for the other lipids between leaflets. Gangliosides DXG1 and DXG3 were added only to the exofacial layer (Fig. 5B).

For the protein components, we designed partial structures to represent the GABA_B_R and the KCC2 transporter. For the receptor component itself, the structure consisted of:

a. the human seven-transmembrane domain (^7TM^B2; blue) of the GABA_B2_ subunit (UniProt accession number: O75899), for which we retrieved the GABA_B2_ 7TM homology model 2, as described by Freyd *et al.* (2017) (87);
b. the coiled-coil domains (cc) within the C-termini of both human GABA_B1_ (B1cc; red) and GABA_B2_ (B2cc; blue) subunits, for which we reconstituted the available x-ray crystal structure (PDB ID: 4PAS) (88);
c. For H8 (green), 23 amino acids from the human GABA_B2_ subunit (sequence ^756^QNRRFQFTQNQKKEDSKTSTSVT^778^; UniProt accession number: O75899) were software-fitted to form an α-helix. H8 is, in fact, a conserved motif among the GPCR superfamily. In the GABA_B2_ subunit, this helical structure connects the 7TM domain to the coiled-coil tail and is exposed to the cytoplasm. Of notice, the H8, 7TM, and extracellular Venus Flytrap (VFT) domains of GABA_B1_, as well as the GABA_B2_ VFT, were not included in our simulations (Fig. 5A).

For KCC2, the native structure consists of a 12-transmembrane helix domain (12TM) flanked on both ends by the cytoplasm-oriented amino and carboxyl termini (63, 89, 90). Based on Wright *et al.* (2017) findings, the 12TM domain is necessary for the association with GABA_B_R, but not the N- and C-termini of the transporter (39). Therefore, we used the 12TM of the bacterial glutamate-GABA antiporter (GadC; PDB ID: 4DJK), which is structurally homologous to the KCC2 transmembrane domain, as previously stated by Agez *et al.* (2017) (63). Here, we refer to this structure as KCC2-*like* (yellow; Fig. 5A).

The previously described PM systems, each coupled with the two partial protein structures, were simulated for 5 µs. For the analyses, we compared time courses of the coordination function, timing for protein-protein and protein-lipid interactions, and major and transient conformational states for a total simulation time of 15 µs.

### Coordination among protein domains varies substantially as membrane cholesterol concentrations are modified

Coordination was the parameter used to quantify dynamic contacts between protein domains, for which we considered both intra- and intermolecular bonds within a specified radius (∼3Å, as deduced from Equation 1, in *Materials and Methods*), through the span of each 5 µs simulation. We thoroughly analyzed the coordination between protein domains over time, and we compared the results with the conformational changes observed in the simulation videos (Videos S1, S2 and S3). This allowed us to identify the major configurational states that protein domains underwent when embedded in different lipid environments, which are summarized in Fig. 6. In addition, time courses of the coordination numbers for protein domain pairs are displayed in Figure S6 (Fig. S6).

**Figure 6.**
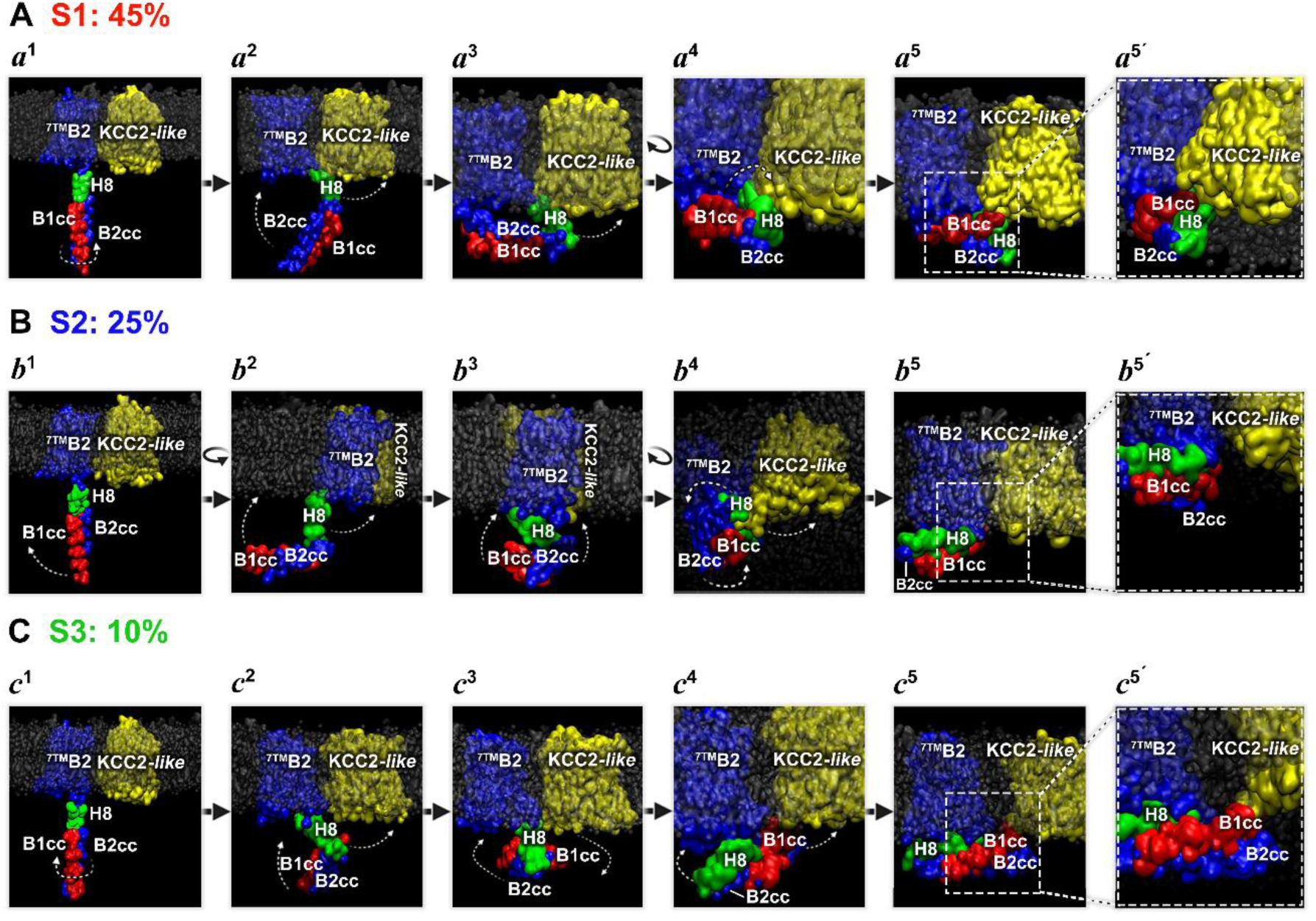
Conformational states for the GABA_B_R tail domains and GABA_B_R:KCC2-*like* interactions are influenced by membrane lipid composition. Although the three assayed system models shared the initial configuration (a^1^, b^1^ and c^1^), the protein components showed highly dynamic and differential behaviors throughout the different simulation trajectories. In A-C, selections of significant protein events exhibit the main differences observed among the membrane setups. **A)** In system 1 (S1: 45%), representative sequences a^1^-a^5^ evince a rotation of the coiled-coil (cc) domains (a^1^), followed by the movement towards the membrane of the H8 motif, which acts as a hinge (a^2^) and brings the B2cc domain closer to the inner leaflet (a^2^ and a^3^). As the GABA_B_R tail folds (a^4^), another rotation of the cc domains allows a stable interaction between B1cc and the KCC2*-like* structure (a^5^ and a^5’^). **B)** In system 2 (S2: 25%), the cc domains move towards the membrane without prior rotation (b^1^) and stay parallel to the lipid bilayer, establishing a 90-degree angle between them and H8 (b^2^). B1cc and B2cc transiently move and finally interact with the lipid components when H8 hinges towards the membrane (b^2^ and b^3^). Once the tail folds, H8 and B1cc interact briefly with KCC2*-like* (b^4^) before moving away (b^5^). The GABA_B_R tail domains remain overlapped and collapsed towards the membrane for the rest of the simulation, but no other contact was detected with the KCC2*-like* structure (b^5’^). **C)** Sequence in system 3 (S3: 10%) exhibits a tail rotation as in S1 (a1), but in the opposite direction (c^1^). Once again, H8 acts as a hinge and brings the cc domains towards the membrane (c^2^). In doing so, B1cc faces the lipid bilayer, yet B2cc remains exposed to the cytoplasm-like space (c^3^). H8 serves as an axis for the horizontal rotation of the GABA_B_R tail that moves towards the KCC2*-like* skeleton (c^3^), granting the B1cc:KCC2*-like* interaction (c^4^ and c^5^). The tail continues to move in a constant and vibrating way, stabilizing this protein-protein contact (c^5’^).

We started with the transmembrane domains ^7TM^B2 and KCC2*-like,* for which we detected substantial differences among lipid bilayer systems (Fig. S6A). For the 45% membrane [CHOL] (S1: 45%), the 7TM and 12TM domains remained apart from each other. However, when cholesterol levels were reduced to 25% (S2: 25%) and then to 10% (S3: 10%), the two TM structures moved closer to each other, which allowed for partial and weak transmembrane interactions. Then, as we descended the GABA_B2_ structure, we investigated the coordination of H8 with KCC2*-like* and the coiled-coil domains. In general, we observed significant differential arrangements, even though the same protein configuration was selected to start every simulation for systems 1-3 (Fig. 6A a^1^, 6B b^1^ and 6C c^1^). However, as simulations proceeded, two shared features were detected among the protein configurations: a) a retraction of the GABA_B_R tail (Fig. 6A a^2-5^, 6B b^2-5^ and 6C c^2-5^), and b) the movement of the H8 domain towards the membrane, at the point where it laid parallel to the PM axis (Fig. 6A a^5-5’^, 6B b^5-5’^ and 6C c^5-5’^). In each system, transient conformational changes observed prior and after these two events were significantly different. For S1: 45% and S3: 10%, a rotation of the cc domains was observed before the H8 helix shifted towards the membrane, with the spin taking opposite directions in each of these two lipid bilayer models (white curved dashed arrows; Fig. 6A a^1^ and 6C c^1^). S2: 25% contrasted with both S1 and S3, in that its cc domains initiated the route towards the PM without any previous rotation at all (Fig. 6B b^2^). Although, they did not reach their destination until H8 hinging momentum dragged them along on its way to the membrane (Fig. 6B b^3^). Then, H8 established contact with KCC2*-like* (Figs. 6B b^4^ and S6B), while strongly interacting with B1cc (Fig. S6C). This differential arrangement appeared to change the orientation of the tail fold, rotating the curled H8, B1cc and B2cc domains in a counterclockwise direction (Fig. 6B b^4-5^) and, therefore, moving the GABA_B_R coiled-coils away from the KCC2*-like* transporter (Fig. 6B b^5-5’^).

We continued to investigate the coiled-coil interactions. The GABA_B1_-GABA_B2_ association is highly stable and resists the action of some detergents, as we previously showed in Figs. 1 and 3. This scenario was also the case *in silico.* In fact, throughout the time course, our simulations showed that the B1cc:B2cc coordination always displayed an oscillatory pattern within a 0.8-1.3 range, for the three membrane systems under study (Fig. S6E). However, the interactions of each cc motif with other protein domains did differ substantially among S1: 45%, S2: 25%, and S3: 10% (Fig. S6C, S6D and S6F). Of particular interest, we detected a direct contact between the B1cc domain and the KCC2*-like* transporter in two out of the three CHOL:PL systems assayed, with remarkable differences in kinetics between them (Figs. 6A a^5-5’^, 6C c^4-5’^ and S6F). For S1: 45% and S3: 10%, B1cc and KCC2*-like* established stable interactions that lasted for the remaining time of simulation, although each protein domain continued to oscillate while in contact (Fig. S6F). For high [CHOL], B1cc:KCC2*-like* coordination took more than 1 µs to become detectable, and it increased following an oscillatory pattern. For low [CHOL], in contrast, the interaction commenced almost immediately from the starting point, and maintained higher coordination throughout, as compared to that observed for S1. For intermediate [CHOL], finally and unlike in the other two systems, B1cc and KCC2*-like* only transiently interacted, and this was due to a brief contact between the 12TM domain and H8 (Figs. 6B b^4^ and S6B). However, immediately after initiating contact, and as previously described for H8, a new series of conformational states in S2: 25% stabilized the GABA_B_R tail away from the transporter, which ended H8:KCC2*-like* interaction (Fig. 6B b^5-5’^).

### Changes in membrane lipids differentially affect the dynamics of protein domains, even those not immersed in the lipid bilayer

Because we had observed that the coordination plots for systems 1, 2, and 3, displayed differential outlooks for any given pair of protein domains analyzed (Fig. S6), our results suggested a substantial influence of PM lipid variations upon protein behavior. We noticed that H8, B1cc, and B2cc all seemed to respond distinctly to membrane cholesterol fluctuations, even though they were not immersed in the computational PM, but instead they were oriented towards the virtual intracellular space. Therefore, to gain a deeper understanding of the effects of lipid variations upon protein dynamics, we analyzed protein-lipid interactions for each membrane system model. First, we investigated coordination over time between the GABA_B_R tail domains and the lipids of the inner leaflet. Surprisingly, we discovered that each protein domain interacted differentially with every cytofacial lipid type. Moreover, the nature of those interactions appeared highly dependent on the global lipid composition (Fig. 7). For instance, H8 mainly interacted with DPPC, DPPE, and DPPS, in the three membrane systems (Fig. 7A a^1^). However, the α-helix 8 coordination with DPPC increased as membrane [CHOL] dropped from 45% to 10%, while by contrast H8 coordination with DPPS decreased as [CHOL] descended (Table S1). On the other hand, the lowest average coordination for H8:DPPE was detected for S3: 10% (Table S1). For S1: 45% and S2: 25%, H8:DPPE coordination values exhibited no substantial differences (Table S1). Overall interactions between the H8 and CHOL molecules were minor, and they were mainly detected in system 2 where intermediate [CHOL] were present in the computational membrane (Fig. 7A a^1^ and 7B b^1^).

**Figure 7.**
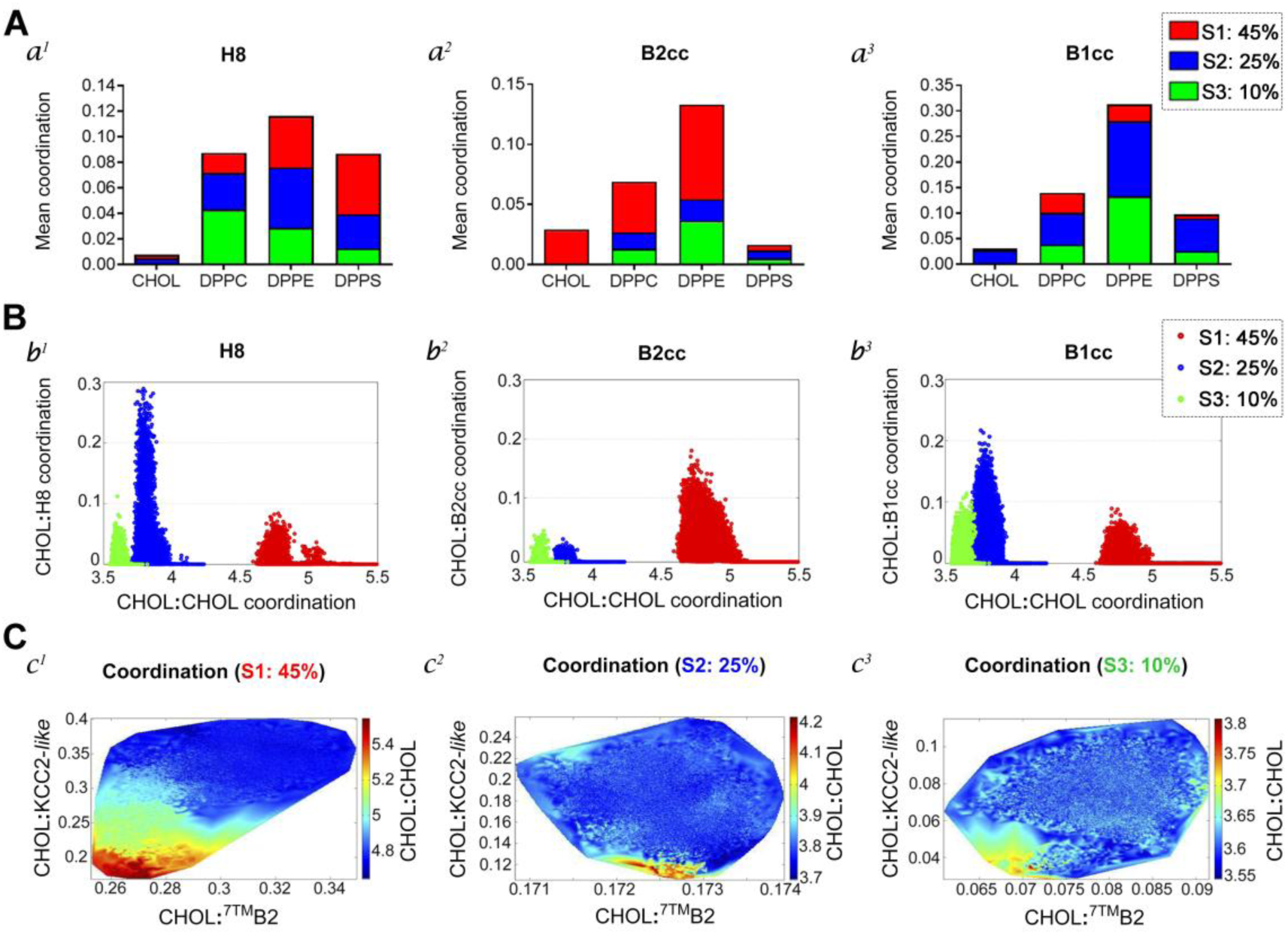
Protein domains interact differentially with cholesterol and other plasma membrane lipids. **A)** Average coordination values among GABA_B_R tail domains: H8 (a^1^), B2cc (a^2^), and B1cc (a^3^), and lipids (CHOL, DPPC, DPPE, and DPPS) from the inner leaflet of the computational plasma membrane, are represented in stacked bar charts. The colors red, blue, and green indicate systems 1 (45% CHOL), 2 (25% CHOL), and 3 (10% CHOL), respectively. **B)** Scatter plots compare coordination between cytofacial cholesterol (CHOL) and GABA_B_R tail domains: H8 (b^1^), B2cc (b^2^), and B1cc (b^3^) (vertical axis), and coordination among CHOL molecules (horizontal axis). Data dispersion for each simulated membrane system model is displayed in colors, as used in A. **C)** Charts compare coordination numbers between global cholesterol and the transmembrane domains of KCC2*-like* (left vertical axis) and GABA_B_R (^7TM^B2; horizontal axis), for systems 1 (c^1^), 2 (c^2^), and 3 (c^3^). Coordination among cholesterol molecules is represented in a heatmap color scale (right vertical axis).

For the B2cc domain, lipid preferences were CHOL, DPPC, and DPPE, and were identified mostly in S1: 45% (Fig 7A a^2^ and 7B b^2^). Relevant B2cc:CHOL interactions occurred only under the highest membrane catechol concentrations. Also, the highest B2cc:DPPC coordination levels were detected for this system. Interestingly, the average coordination for B2cc:DPPE in S1: 45% was more than double the one detected in system 3 (Table S1), but the lowest coordination values for this pair were observed in system 2, with 25% CHOL (Table S1).

The B1cc domain also interacted preferentially with DPPE (Fig. 7A a^3^), with elevated coordination levels for systems 2 and 3 (Table S1). Furthermore, the average coordination values detected in S2: 25% for the pairs B1cc:DPPC, B1cc:DPPE, and B1cc:DPPS, were considerably higher than the ones observed for H8 and B2cc and any other lipid, for all three systems that were analyzed (Table S1).

Based on our observations that transmembrane domain interactions were somehow influenced by membrane lipid composition in a way that contacts could either occurred or not (Fig. S6A), we further compared coordination between TMs and cholesterol molecules to gain a deeper mechanistic insight into the interplay between lipids and proteins. For this analysis, coordination was no longer defined as an intermittent parameter resulting from fleeting associations, such as those of the GABA_B_R tail domains as have been described herein. Instead, we considered coordination as a continuous variable of the TM domains as they remained within the limits of the lipid bilayer and constantly interacted with all its lipid components. Our findings are displayed in heatmaps in Fig. 7C. For S1: 45%, we found that the highest cholesterol self-coordination occurred when the lowest CHOL:TM coordination values were observed (Fig. 7C c^1^). Hence, stronger CHOL:CHOL interactions took place when the catechol molecules were not in proximity to the TM structures. As membrane [CHOL] were reduced to 25% and then to 10%, significant CHOL:CHOL interactions appeared at systematically lower values of coordination for both CHOL:KCC2*-like* and CHOL:^7TM^B2 (Fig. 7C c^2-3^). In these two membrane system models, the highest levels of cholesterol self-coordination seemed to require minimal CHOL:KCC2*-like* interactions, concomitantly with a progressive reduction in CHOL:^7TM^B2 coordination. Although, some minimal level of CHOL:^7TM^B2 coordination was still necessary.

Based on the observation that CHOL molecules interacted with differential affinity with each of the transmembrane proteins, and that they apparently preferred the ^7TM^B2 segment more than the 12TM domain of the KCC2*-like* structure, we investigated the ^7TM^B2 sequence for potential hotspots for cholesterol interactions (Fig. S7). We used the Fuzzpro tool from the EMBOSS software to identify the two putative consensus linear motifs, CRAC [N-terminal-(L/V)-X_1–5_-(Y)-X_1–5_-(K/R)-C-Terminal] and CARC [N-terminal-(K/R)-X_1–5_-(Y/F/W)-X_1–5_-(L/V)-C-terminal], along the human GABA_B2_ sequence (UniProt accession number: O75899) (91, 92). We identified one CRAC motif in the fifth transmembrane domain (TM5), whose sequence was ^657^LGIVYAYK^664^. In addition, we recognized three other regions that matched CARC consensus motifs: a) the first one: ^476^KISLPLYSILSAL^488^, was identified in TM1, located immediately in the membrane insertion from the extracellular space; b) the second one, whose sequence was ^556^RTWILTV^562^, was found in TM3, and c) the third sequence: ^664^KGLLMLFGCFL^674^, was in TM5. Following identification, we measured coordination of all the CRAC and CARC motif sequences with respect to CHOL for systems 1-3 (Fig. S7). Overall, we discovered that the occupancy of each interaction site varied according to global membrane cholesterol concentration. In general, lower coordination numbers were detected for all four sequences in S3: 10%, whereas the highest interactions were found in S1: 45%. In system 1, we observed that the CRAC and CARC motifs identified in TM5 maintained coordination numbers between 0.5-1.5 with CHOL, and sometimes even higher. These interaction hotspots seemed to lose strength as membrane cholesterol concentration decreased. Nevertheless, cholesterol molecules were stabilized within the coordination radius for these two ^7TM^B2 sequences throughout the simulation, in the three membrane systems that were assayed. This was true despite any changes in membrane [CHOL] levels. On the other hand, the coordination numbers between the CARC motif in TM1 and CHOL oscillated around 0.5 in systems 1 and 2, but levels for this pair substantially decreased when cholesterol concentrations reached 10% (S3: 10%). Finally, the CARC sequence found in TM3 only became significant throughout the simulation of S1: 45%, with coordination numbers around 0.5.

### For systems 1 and 3, B1cc:KCC2*-like* interactions occur not only at different times, but also involve distinct amino acid residues

In consideration of our observation that B1cc:KCC2*-like* interactions took place at different times during the 5 µs simulations for systems 1 (S1: 45 %) and 3 (S3: 10%) (Fig. S6F), and that highly distinct conformational states were observed in both systems prior to achieving the protein-protein contacts (Fig. 6A and 6C), we performed a more exhaustive comparative analysis at the amino acid level, to investigate which residues were involved in the these two computational PM models. In S1: 45%, the interaction was identified between residues located in the N-terminal region of B1cc: ^6^EEEKSRLLEK^15^ corresponding to ^884^EEEKSRLLEK^893^ in the human GABA_B1_ subunit (UniProt Q9UBS5), and a sequence present in the intracellular loop 4 (ICL4) of the KCC2*-like* structure, comprising residues 325-335 (Fig. 8A). Interestingly, among the B1cc amino acids that were involved, we found a di-leucine internalization signal: ^886^EKSRLL^891^ (numbering according to the human GABA_B1_ subunit). This consensus motif is known to play a crucial role in regulating the availability of the GABA_B_ receptor on the cell surface (93–95). In contrast to system 1, the contact surface in S3: 10% was comprised of entirely different residues in both B1cc and KCC2*-like* domains. For the antiporter, we identified the interacting area to be defined among amino acids 140 and 151, involving the intracellular loop 2 (ICL2; Fig. 8B). Moreover, when membrane cholesterol concentrations were dropped to 10%, we found that the di-leucine internalization motif within B1cc no longer participated in the B1cc:KCC2*-like* interaction. Instead, a slightly wider contact surface was detected in S3: 10%, whose sequence ^31^VSELRHQLQS^40^ (corresponding to ^909^VSELRHQLQS^918^ in the human GABA_B1_ subunit) matched to a region of the B1cc C-terminus.

**Figure 8.**
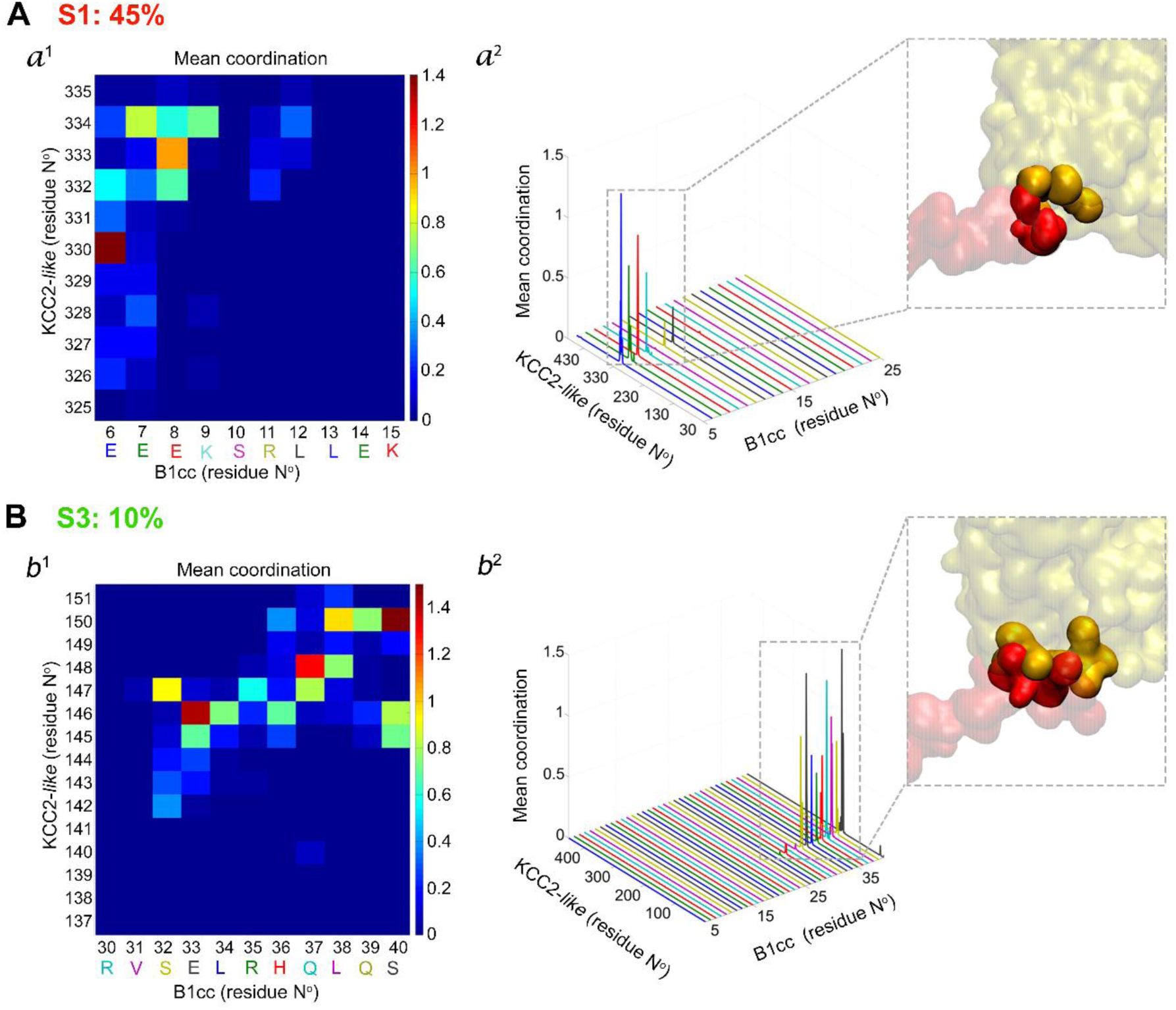
Amino acids residues involved in the B1cc:KCC2*-like* interactions for systems 1 and 3. Average coordination between B1cc and KCC2*-like* at the main contact points in S1: 45% and S3: 10%. Out of several transient associations, only those that prevailed over simulation time are displayed in A and B. **A)** For system 1, heatmap (a^1^) exhibits coordination range for individual amino acids that were recognized within the B1cc:KCC2*-like* contact region. The residues were numbered according to PDB ID: 4PAS and PDB ID: 4FJK sequences, respectively. Amino acids within the B1cc N-terminus are displayed in colors that match mean coordination peaks exhibited in a separate distribution chart (a^2^). The inset features both protein domain arrangements and interacting residues represented according to the Martini coarse-grained (CG) model, where B1cc is shown in red and KCC2*-like* is represented in yellow. **B)** In system 3, the same analyses revealed a slightly wider contact region located at the C-terminus of B1cc. As for the antiporter, the amino acid sequence of B1cc involved in the protein-protein interaction was also different from the one identified in system 1. b^1^ and b^2^ show the implicated residues and their levels of coordination. In b^2^, chart features mean coordination peaks between B1cc and KCC2*-like* residues in the protein-protein interface, while the inset shows the Martini representations of protein domains and specific interacting residues.

## Discussion

Correct brain function requires that neuronal circuits precisely balance excitatory and inhibitory responses (96, 97). However, an excitatory versus inhibitory imbalance has been associated with brain aging and neurodegenerative processes (2, 11, 21, 98, 99). Because of the age-associated brain decline, membrane lipid metabolism can also suffer alterations (3, 4). In the aging cerebellum, heterogeneous lipid level fluctuations and changes in inhibitory signaling have been described (5, 13, 14, 17–21, 75). Although, little is known about the impact of membrane lipid variations on cell surface protein organization, protein-protein interactions, and downstream signaling (8). Therefore, we have investigated how aging impacts the cerebellum, focusing on the protein-protein and protein-lipid interplays between members of the GABAergic system, particularly the GABA_B_ receptor and the KCC2 transporter, and membrane lipids, especially cholesterol.

First, we studied protein expression and distribution of the GABA_B_R subunits along with KCC2. Most notably, we discovered a significant drop in GABA_B2_ levels in the aged rat cerebella, as compared to cerebellar samples from younger rodents. Further immunohistochemical analysis revealed that the age-related GABA_B2_ decrease followed a gradient in the molecular layer (ML). The GABA_B2_ signal appeared to decay gradually from P3m to P18m, and then became barely detectable at P21m. Densities of GABA_B1_ and GABA_B2_ subunits result heterogenous along the different lobules of the rat cerebellar cortex (50, 55). However, we detected a substantial reduction in the global GABA_B2_ expression that extended to all lobulated areas of the aged tissue. Previous studies have reported variations in protein expression levels for GABA_B_R subunits, although the nature of those changes seems to rely on sex, age, species, and the brain region under study (98–100). Deficiency in GABA_B2_ levels can dramatically affect morphological maturation, growth, and migration of immature cortical neurons, as subunit knockdown experiments have shown (101). However, the possible impact of a progressive GABA_B2_ downregulation on fully differentiated and mature neurons is, to the best of our knowledge, unknown. In contrast to GABA_B2_, GABA_B1_ levels and overall distribution did not exhibit significant changes with respect to aging. Functional GABA_B_Rs only reach the cell surface as heterodimers (64, 102–105). Therefore, as the decay of GABA_B2_ proceeds over time, it could ultimately affect the membrane availability of the receptor and even impair downstream signaling, in aged rats. To further characterize the presence of GABA_B_Rs and KCC2 in the aged cerebellum, we applied tandem mass spectrometry and found peptides that were consistent with GABA_B1a_, GABA_B1b_, GABA_B1c_, GABA_B1d_, GABA_B1f_, GABA_B2_, KCC2a, and KCC2b. All the GABA_B1_ isoforms, named GABA_B1a-g_, originate from the same locus but they either undergo mechanisms of alternative splicing or they rely on different promoters and transcription initiation sites (32–35, 106–109). GABA_B1e_ and GABA_B1g_ were not included in our analysis because we focused on analyzing isoforms that could potentially participate in transforming inhibitory circuits as aging progresses. However, both isoforms, GABA_B1e_ and GABA_B1g_ do play other roles. GABA_B1e_ has been described as a signaling-impairing isoform that arrests GABA_B2_ when paired together (106). GABA_B1g_ expression has been described as developmentally regulated, with concentrations dropping acutely after postnatal day 4 (107). The KCC2 transporter can either be homodimerized or heterodimerized, in order to form stable and SDS-resistant forms (41, 63). Our combined WB and UHPLC-MS/MS results proved the presence of KCC2 dimers in both young adult and aged cerebella. Differences in expression of dimeric and monomeric forms were detected but overall KCC2 levels did not exhibit substantial variations with age.

GABA_B_Rs and KCC2 interact to a certain degree in different brain regions (38, 39). Moreover, GABA_B_R activation can induce the internalization of the transporter, which in turn increases the intracellular chloride concentration (39). Ultimately, these actions can lead to a shift in the driving force for the ionotropic GABA_A_R (39). In the cerebellum, a finely modulated balance between GABA_A_R and GABA_B_R actions regulates neuronal activity and synaptic plasticity (54, 56, 110, 111). Our co-immunoprecipitation reactions confirmed that a fraction of the GABA_B_R pool interacts with KCC2 and GABA_A_Rs, establishing a multiprotein complex. However, when comparing GABA_B_R to KCC2 association levels, we discovered that they were influenced by age or any age-related process affecting cerebellum homeostasis. All previously mentioned protein interactions could either involve direct physical contact or require the participation of additional partners to ensure the receptor-transporter coupling. In either case, such close-distance arrangement of proteins suggests that an intricately cooperative system likely exists between the ionotropic and metabotropic GABA receptors (112, 113). For cerebellar Purkinje cells (PkC), at parallel fiber to PkC and interneurons to PkC synapses, this crosstalk mechanism would be strategically important to ensure synaptic plasticity throughout well-integrated and tightly tuned responses (22, 56, 58, 111, 114–116).

As age progresses, cells naturally and progressively decline, in tune with multiple architectural changes (9). The presence of lipofuscin (LP) aggregates is considered a confirmatory and distinctive feature of aging (9, 117). LP deposits are intracellular storage bodies that accumulate progressively over time as the cellular ability to degrade residual materials decreases (9, 118). In our analysis, the comparison between P18m and P21m cerebellar sections revealed a progressive buildup of LP aggregates within the cytoplasm of PkCs. Interestingly, in P21m, extracellular LP accumulation also became more evident around the PkC somata, which could be related to increased apoptosis (9, 118). Age-related enlargement of LP deposits can also result from a hampered lipid and protein metabolism (9). After finding changes in protein-protein interactions in the aged cerebellum, along with significant LP deposits, we further analyzed lipid composition profiles to assess whether they were also affected over time. Multi-domain neurotransmitter receptors, such as GABA_B_Rs and GABA_A_Rs, usually organize themselves into plasma membrane lipid rafts enriched in cholesterol and sphingolipids (46, 49, 119). Such specialized lipid microdomains could play a critical role in modulating protein dynamics and downstream signaling (119–122). Our comparative analysis between young and aged lipid extracts revealed substantially increased levels of cholesterol and sphingomyelin in the older cerebella. Fluctuations in concentrations of cholesterol, gangliosides, sphingolipids, and ceramides, have been associated with LP accumulation, aging, and aging-related neurodegenerative processes (4, 6, 9, 13, 75–80, 123, 124). However, up until now, lipid trends have mostly been determined by the animal model used, sex, age, brain region under analysis, and cell types (2, 4, 5, 13, 14, 75, 76, 80). Considering that the cerebellum is not a specialized repository for lipid storage, our results imply a direct correlation between global lipid variations and plasma membrane lipid modifications that may occur as the cerebellum ages.

Furthermore, we used molecular dynamics (MD) simulations to reach a greater level of understanding of the protein-protein and protein-lipid interplays between GABA_B_Rs and the transporter (125). Recently, structural models for GABA_B_R, in both its active and inactive states, were made available (126–129). For our simulations, however, we used a simplified partial GABA_B_R, along with an additional transmembrane protein whose structure resembled that of KCC2. These simulations were conducted using three lipid bilayer systems (S) with symmetrical distribution of specific cholesterol concentrations: 45% in S1, 25% in S2, and 10% in S3. Generally, we found that preferences for specific lipid types, protein-protein and protein-lipid coordination values, conformational states, and timing, were exclusive for each protein domain, and they differed substantially depending on the global lipid composition of the computational plasma membrane simulated. Parameters such as membrane thickness, lipid headgroups, acyl tail lengths, saturation, glycosylation, and cholesterol content, have been shown to influence protein structure, stability, conformational dynamics, and clustering with other integral membrane proteins (122, 125, 130–136). Hence, our results highlight the mutual influence that lipids and protein domains exert on each other. MD simulations also unveiled a direct interaction between the GABA_B1_ tail (B1cc) and the transporter that occurred in S1: 45% and S3: 10%. Before these associations, however, protein domains displayed different intermediate conformational states in both systems. In S2, no such interaction was detected. Conversely, B1cc adopted a differential configuration that allowed substantial coupling with lipids of the inner leaflet of the computational bilayer. This alternative conformation probably accounts for the lack of association with the KCC2*-like* structure in S2: 25%. In all three systems, the H8 domain played a pivotal role in driving all conformational movements executed by the coiled-coil tails, just as a neck would direct the head’s motion. H8 is a conserved amphipathic helix in class A and class C GPCRs (120). This domain usually adopts a location parallel to the plasma membrane, forming a polar-hydrophobic interface between the GPCŔs tail residues and the lipids from the inner side of the cell bilayer (120, 137). Although it is not immersed in the lipid environment, H8 interacted with the membrane in all simulations, influencing perhaps further interactions between B1cc and B2cc domains and the internal lipids.

We also analyzed amino acid residues involved in the B1cc-transporter associations. Here, we discovered that a di-leucine internalization motif within B1cc itself was involved in the interaction that we observed in S1: 45%. This sequence is usually buried within the GABA_B1_-GABA_B2_ heterodimeric interface and is essential for regulating the receptoŕs cell surface availability (88, 93–95, 138). When this signal is unmasked, GABA_B_Rs undergo clathrin-dependent internalization, while still preserving their dimeric assembly (94, 95). KCC2 is also constitutively internalized via clathrin-mediated endocytosis and binding of the adaptor protein-2 (AP-2) to a non-canonical di-leucine motif present in the C-terminus of the transporter (139). As mentioned before, upon activation, GABA_B_Rs are capable of regulating their own surface levels, and KCC2’s surface levels also (39). Therefore, perhaps the exposure of di-leucine internalization motifs might function as a trigger signal that orchestrates the combined endocytosis of GABA_B_R and KCC2. Since the B1cc di-leucine motif was only involved in the receptor-transporter association observed in S1: 45%, and not in the other two system models (S2: 25% and S3: 10%), then we speculate that changes in membrane lipid composition, such as those observed in aging and in neurodegenerative processes, might compromise GABA_B_R-KCC2 interplay and their influence in GABA_A_R functionality. Finally, we focused on the GABA_B2_ transmembrane domains (^7TM^B2) since they play a pivotal role in regulating GABA_B_R transactivation and G-protein mediated signaling (64, 127, 129, 140). Cholesterol molecules have been identified within the GABA_B_R heterodimeric structure (128, 129). They seem to reside specifically inside the interfacial space delimited by the GABA_B1_ and GABA_B2_ TM5 domains, which serves to stabilize the receptor inactive state (128, 129). We explored how cholesterol interacted with ^7TM^B2 in S1: 45%, S2: 25% and S3: 10%, focusing on *in silico-*identified cholesterol recognition hotspots within the ^7TM^B2 sequence, named CRAC and CARC (91). Importantly, we uncovered two motifs, one CRAC and one CARC, within the ^7TM^B2 TM5. The coordination numbers between these individual CHOL recognition sites and CHOL molecules revealed substantially different behaviors under each membrane system that we analyzed. CHOL recognition motifs are believed to exert a crucial role in incorporating GPCRs into lipid rafts (141). Therefore, fluctuations in membrane lipid levels, such as those described herein as being related to aging, could undermine GABA_B_R signaling by potentially affecting the receptor recruitment into membrane lipid domains or by altering the interfacial TM domain arrangements that stabilize the RGABA_A_-RGABA_B_ heterodimer.

Overall, the presented results suggest that regulation of membrane protein dynamics is highly specific and sensitive to fluctuations in global lipid composition, such as those widely described for aging and neurodegenerative processes (8, 125, 136, 142). Besides cerebellar motor functions, the cerebellum receives and delivers multiple inputs and responses that integrate with other brain areas (22, 143, 144). Therefore, variations in cerebellar circuits could ultimately affect movement, balance, attention, perception, learning, language, and even emotional processing in the elderly (22). Although our results might only have scratched the surface, we hope to inspire new research efforts to investigate further membrane regulation and GABAergic dynamics in aging and age-related pathological circumstances.

## Materials and Methods

### 1 Animals

All procedures were performed according to guidelines established by the U.S. National Institutes of Health’s Guide for Care and Use of Laboratory Animals, the Animal Research: Reporting in vivo Experiments (ARRIVE) Guidelines, and the Institutional Animal Care and Use Committee (IACUC), School of Medicine, National University of Cuyo, Mendoza, Argentina (Protocol IDs: 74/2016; 141/2018; 152/2019; 218/2022). Male Wistar rats were maintained in standard housing conditions for 3, 18 or 21 months (P3m, P18m, and P21m, respectively), under a 12:12 light-dark cycle, and unlimited access to water and food. Animals were briefly stunned using CO_2_ and sacrificed by decapitation. For immunohistochemistry (IHC), cerebella were removed and immediately immersed in 4% paraformaldehyde in phosphate-buffered saline (PBS, pH 7.3) for overnight fixation. For Western blot (WB) and co-immunoprecipitation (Co-IP), cerebella were snap-frozen in liquid nitrogen and stored at -80°C until further processing.

### 2 Western blot

GABA_B1_, GABA_B2_ and KCC2 protein levels in young adult and aged rat cerebella were assayed via Western blot (WB), as previously described (145, 146). Briefly, total protein extracts were obtained from 60 mg of P3m and P18m rat cerebella (7 rats per age). For each sample, a double-extraction procedure was performed with 200 µL of Triton X-100 lysis buffer supplemented with protease and phosphatase inhibitors [LB: 50 mM Tris-HCl pH 7.5, 150 mM NaCl, 5 mM EDTA, 0.5% (v/v) Triton X-100, 10 mM NaF, 10 mM Na_3_VO_4_, 1 mM PMSF and 1X solution of Complete Mini Protease Inhibitor Cocktail (Roche Applied Science, Mannheim, Germany)], in order to increase the plasma membrane (PM) protein yield. Protein concentrations were quantified using the Pierce^TM^ BCA Protein Assay Kit (Pierce Biotechnology, Thermo Fisher Scientific Inc., Waltham, MA, USA). For each sample, 10-30 μg of proteins in Laemmli buffer with 10% β-mercaptoethanol were denatured at 70°C for 5 minutes, then cooled down to room temperature, and resolved in 8% SDS-PAGE gels. After semi-dry electroblotting, PVDF membranes were incubated with blocking solution [5% (w/v) skim milk in either Tris-buffered saline (TBS) or PBS with 0.1% (v/v) Tween-20]. Then, they were rinsed and incubated overnight (4°C) with the primary antibody solution (for further information see the antibody section). Subsequently, membranes were washed and incubated with the secondary antibody conjugated with horseradish peroxidase (see antibody section). Proteins reacted with Immobilon® Western Chemiluminescent HRP Substrate (EMD Millipore, Burlington, MA, USA), and bands were detected with ImageQuant^TM^ LAS4000 (GE Healthcare Life Sciences, Pittsburgh, PA, USA). Band intensities were normalized to total protein per lane by using the Coomassie Brilliant Blue R-250 (CBB R-250) staining method, as previously described (145–147). By using Image Lab Software^TM^ 6.0.1 (Bio-Rad Laboratories Inc.), the blot was merged with the CBB-stained membrane, then the background was subtracted out, and the lanes and bands were defined. Each band of interest was normalized to the total CBB R-250 intensity in its corresponding lane. Densitometric values were compared using Graph Pad Prism 9.0.2 (GraphPad Software Inc.).

### 3 Co-immunoprecipitation

Cerebellar tissue (∼150 mg) was homogenized in NP-40 lysis buffer supplemented with protease and phosphatase inhibitors [50 mM Tris-HCl pH 7.5, 75 mM NaCl, 5 mM EDTA, 0.05% NP-40, 2% sodium deoxycholate, 10 mM NaF, 10 mM Na3VO4, 1 mM PMSF and 1X solution of Complete Mini Protease Inhibitor Cocktail (Roche Applied Science)], using the two-step procedure previously described for WB. After estimating total protein concentrations, extracts were diluted to a final concentration of 1µg/µl using wash buffer (50 mM Tris-HCl pH 7.5, 75 mM NaCl, 5 mM EDTA, 10 mM NaF, 10 mM Na_3_VO_4_, 1 mM PMSF and 1X solution of Complete Mini Protease Inhibitor Cocktail). Precleared samples were incubated with guinea pig anti-GABA_B2_, mouse anti-GABA_B1_, and rabbit anti-KCC2 antibodies (see antibody section) for two hours in constant agitation at 4°C. Then, previously washed, and stabilized Protein A/G agarose beads (sc-2003, Santa Cruz Biotechnology Inc., Dallas, TX, USA) were added to the samples at a bead/tissue extract ratio of 1 µl/3.5 µl. Subsequently, the tubes were incubated in agitation overnight at 4°C. After the incubation period, the samples underwent centrifugation, and the resulting supernatant (Sup), representing the protein fraction not bound to the immunocomplexes, was separated from the precipitate, and stored in sample buffer at -20°C. The obtained precipitates were rinsed twice in wash buffer, followed by centrifugation. After these two purification cycles, the immunocomplexes (IP) were collected. Subsequently, the IPs were eluted in sample buffer, along with the supernatants obtained after each wash step (Wash 1 and Wash 2). Finally, the samples were denatured and resolved by SDS-PAGE in 8% gels, as previously outlined in the WB section.

### 4 Immunohistochemistry

After overnight fixation, tissues were washed twice in PBS, and gradually dehydrated in ethanol of increasing concentrations: 50, 70, 80, 96, and 100%. After these steps, cerebella were rinsed twice in xylene, immersed in a 1:1 xylene/Histoplast mixture, and finally included in Histoplast (Biopack, Buenos Aires, Argentina), as previously described (145, 146, 148–151). Ten-micrometer paraffin-embedded cerebellar sections were cut, mounted on positively charged slides, and stored at 4°C until further processing. For immunolabeling, slides were gradually rehydrated in decreasing concentrations of ethanol (100, 96, 80, 70, and 50%), and finally washed in MiliQ water. For antigen retrieval, tissue sections were immersed in boiling citrate buffer (10 mM sodium citrate pH 6, 0.05% Tween-20) for 30 minutes, and washed three times in PBS, in order to re-equilibrate pH to physiological conditions (pH 7.2). To prevent non-specific binding, slides were incubated in blocking solution [10% (w/v) normal donkey serum, 1% (v/v) Triton X-100, 0.2% (w/v) gelatin in 1X PBS] for 1 hour at room temperature (RT), inside a humidity chamber. Immediately afterwards, slides were incubated overnight with primary antibodies, then washed with PBS and exposed to secondary antibodies for two hours. Both primary and fluorophore-conjugated secondary antibodies were diluted in dilution buffer [2% (w/v) normal donkey serum, 1% (v/v) Triton X-100, 0.2% (w/v) gelatin in 1X PBS]. After washing with 1X PBS, slides were attached to coverslips using mounting media with the nuclear markers propidium iodide (PI) or DAPI (D1306, Life Technologies-Invitrogen). After the immune reaction, sections were analyzed using a confocal microscope Olympus FluoView FV-1000 (Olympus America Inc., Center Valley, PA, USA). Images were further processed with Image J 1.49v (National Institutes of Health, USA) and edited with Adobe Photoshop CC v2021 (Adobe Systems Inc., San Jose, CA, USA).

### 5 Antibodies

The following primary antibodies were used: rabbit monoclonal anti-GABA_B_ Receptor 2 (Abcam Inc., Waltham, MA, USA; ab75838; RRID: AB_1310245) dil. WB: 1:5000; (IP) WB: 1:500-1:3000; IHC: 1:400; guinea pig polyclonal anti-GABA_B_ Receptor 2 (EMD Millipore, AB2255; RRID: AB_10563515) dil. IP: 1:100; mouse monoclonal anti-GABA_B_ Receptor 1 (Abcam Inc., ab55051; RRID: AB_941703) dil. WB: 1:25000 (72); IP: [2,5 µg]; (IP) WB: 1:500-1:10000; IHC: 1:500; rabbit polyclonal anti-KCC2 (GeneTex, Irvine, CA, USA; GTX133130; RRID: AB_2886837) dil. WB: 1:30000; IP: [5 µg]; (IP) WB: 1:5000-1:10000; IHC: 1:500; rabbit polyclonal anti-GABA_A_ Receptor α1 (EMD Millipore, 06-868; RRID: AB_310272) (IP) WB:1:500-1:1000. Anti-rabbit (Jackson ImmunoResearch Laboratories Inc., West Grove, PA, USA; 711-035-152; RRID: AB_10015282) and anti-mouse (Jackson ImmunoResearch Laboratories Inc., 115-035-003; RRID: AB_10015289) peroxidase-conjugated secondary antibodies generated in donkey were used for WB at 1:50000 dilution. The fluorophore-conjugated secondary antibodies included in this study were Alexa Fluor® 488-congugated anti-rabbit IgG (Jackson ImmunoResearch Laboratories Inc., 711-545-152; RRID: AB_2313584), dil. IHC: 1:1000, and Cy^TM^3-conjugated anti-mouse IgG (Jackson ImmunoResearch Laboratories Inc., 715-165-151; RRID: AB_2315777) dil. IHC: 1:1000.

### 6 Mass spectrometry analysis

#### 6.1 Sample preparation

Total cerebellar protein extracts were generated from aged male Wistar rats (P18m). For protein extraction, two lysis buffers with different detergent compositions and ionic strengths were used to homogenize tissue samples. Protein extracts obtained alternatively with NP-40 and Triton X-100 lysis buffers were seeded in triplicates and resolved in two separate gels by SDS-PAGE. Following electrophoresis, gels were stained with Coomassie Brilliant Blue G250. Then, printed-to-scale immunoblots for KCC2 and GABA_B_ subunits were used as a reference to identify and excise stained gel fragments potentially containing proteins of interest. Bands above 200 kDa, observed in GABA_B_R subunits and KCC2 blots (Fig. 1A), were considered for analysis based on evidence that suggested that both the GABA_B_ receptor and the transporter form homo- and/or heterodimers (41, 63–66). Five gel fragments were obtained from each gel (Fig. 1C). Designated fragment 1 was excised at ∼280 kDa, where a high molecular band was observed for KCC2 (Fig. 1A). Fragment 2 was cut out immediately below fragment 1, at ∼250 kDa, where bands at higher molecular weights were identified in GABA_B1_ and GABA_B2_ blots. At ∼130 kDa, we obtained fragment 3, consistent with the prominent band observed in the KCC2 blot. Fragment 4 was defined to contain the band observed for GABA_B2_, and the two bands observed for GABA_B1_ between the 95 kDa and the 130 kDa molecular weight markers. Finally, fragment 5 was excised right below fragment 4, at ∼95 kDa, where the lower band observed in the GABA_B1_ blot was expected. After excision, gel fragments were digested with trypsin and in-line separated by ultra-high performance liquid chromatography–tandem mass spectrometry (UHPLC-MS/MS). By performing tandem mass spectrometry with a customized database that included the sequences of GABA_B1a_, GABA_B1b_, GABA_B1c_, GABA_B1d_, GABA_B1f_, GABA_B2_, KCC2a, and KCC2b (protein accession numbers: CAA71398.1, CAA71399.1, BAA34708.1, BAA34709.1, AAK69540.1, O88871, Q63633, and Q63633-2), we recovered isoform-specific peptides.

#### 6.2 UHPLC-MS/MS analysis

The excised gel bands were further processed according to Liu *et al.* (2021) and Paez & Callegari (2022) (152, 153). Proteins were in-gel reduced and alkylated with 10 mM DTT (Sigma-Aldrich, Saint Louis, MO, USA) and iodoacetamide (Sigma-Aldrich), respectively, followed by digestion using sequencing grade trypsin (Promega, Madison, WI). As previously described (154–156), the tryptic peptides were in-line desalted through an Acclaim^TM^ PepMap^TM^ 100, 75 µm x 2 cm nanoViper^TM^ C18, 3 µm, 100Å (Thermo Fisher Scientific Inc.), using water/acetonitrile/trifluoroacetic acid (TFA) (98:2:0.05% v/v) at 6 µl/min, during 5 minutes. Then, they were separated by an Acclaim^TM^ PepMap^TM^ 100, 75 µm x 15 cm nanoViper^TM^ C18, 2 µm, 100Å (Thermo Fisher Scientific Inc.), for 110 minutes. The mobile phase (A) consisted of water/formic acid (99.9:0.1% v/v), while phase (B) components were water/acetonitrile/formic acid (20:80:0.08% v/v). Fisher Chemical™ Optima™ grade LC-MS solvents were used (Thermo Fisher Scientific Inc.). The standard gradient was 0-4 min, 1.2% (B) isocratic; 4-98 min, 3-98% (B) linear, followed by equilibration of (A):(B) (98:2%). Solvent flow rate was established at 300 nl/min. The separation was performed using Ultimate 3000RS UHPLC in nanoflow configuration (Thermo Fisher Scientific Inc.), coupled to QExactive Plus quadrupole Orbitrap (Thermo Fisher Scientific Inc.) through a nanoelectrospray ion source, using FullMS followed by ddMS^2^ (DDA) for 110 minutes, as previously described in Kelstrup *et.al* (2012), Sun *et.al.* (2013) and Pizarro-Guajardo *et.al.* (2018) (157–159). In-house licensed Mascot Distiller v2.6.2.0 (www.matrixscience.com, UK) and Proteome Discoverer v2.1 (Thermo Fisher Scientific Inc.) were used to generate the peak list at the Mascot generic format (mgf) from the original raw file. To identify +1 or multiple charged precursor ions from the mass spectrometry data file, parent mass (MS) and fragment mass (MS/MS) peak ranges were 300-1800 Da (resolution 70000) and 65-2000 Da (resolution 17500), respectively.

#### 6.3 Bioinformatics analysis

Mascot server v2.7.1 (www.matrix-science.com) in MS/MS ion search mode (local licenses) was applied to conduct peptide matches (peptide mass and sequence tags) and protein searches against database SwissProt (http://www.uniprot.org) v2020_4 (563082 sequences; 202799066 residues), with taxonomy filter for *Rattus novergicus* (SP-rat) (8120 sequences), as well as in-house customized databases such as All_GABA receptors, and KCCs (data from www.uniprot.org). The following parameters were set for the search: carbamidomethyl (C) on cysteine was set as fixed, and variable modifications included asparagine and glutamine deamidation and methionine oxidation. Only one missed cleavage was allowed; monoisotopic masses were counted; the precursor peptide mass tolerance was set at 20 ppm; fragment mass tolerance was established at 0.02 Da, and the ion score or expected cut-off was set at 5. The MS/MS spectra were searched with Mascot using a 95% confidence interval (C.I.%) threshold (*p* < 0.05), for which minimum scores of 25 and 13 were used for peptide identification for SP-rat and customized databases, respectively. This indicates identity or extensive homology. Peptides unassigned with a protein hit match were selected and used for extra protein identification, previous corroboration with each MS/MS or fragmentation map as well as through BLAST-P (https://www.ncbi.nlm.nih.gov/). Also, the error tolerant mode was modified at Mascot search to corroborate potential peptides that were unidentified after the first search. When the identified peptides matched multiple protein IDs equally well, then only those proteins that appeared in at least two or more replicates were considered valid to be included in the list.

### 7 Lipid analysis

#### 7.1 Lipid extraction

For lipid extraction, P3m (n=4) and P18m (n=5) cerebella were collected and processed following the Bligh & Dyer (1959) method (160). Briefly, tissue samples (∼250 mg) were homogenized in a chloroform/methanol (1:2 v/v) solution using a Bio-Gen PRO200 homogenizer (PRO Scientific Inc., Oxford, CT, USA). The homogenates were stored at -20°C until further processing. One volume of chloroform and 1.8 volumes of water were added, reaching a final 2:2:1.8 (v/v) chloroform/methanol/water ratio. Then, the tubes were vigorously vortexed and centrifuged for 20 minutes at 2000 rpm. After separating the upper methanolic-aqueous phase, the lower chloroform phase containing the lipids was recovered. The interface, containing denatured proteins, was washed twice with the same solvent mixture, and the lower phases were recovered again to be combined with the previously obtained chloroform extract to ensure efficient lipid recovery. The obtained lipid extracts were dried under a stream of N2; later, they were resuspended in known volumes of chloroform/ methanol 2:1 (v/v) and stored at -20°C until use. Specific volumes of the extracts were used to: a) determine the total lipid phosphorus content; b) determine the cholesterol content; and c) separate lipids into different classes using high-performance thin-layer chromatography (HPTLC). HPLC grade solvents were used (Dorwill S.A., Grand Bourg, Buenos Aires, Argentina).

#### 7.2 Determination of total lipid phosphorus

Total phospholipids (PL) recovered from tissues, as well as different PL classes separated by HPTLC, were subjected to phosphorus content determination, following a modified version of the Rouser *et al*. (1970) method (161). This method involves releasing inorganic phosphate from PL through high-temperature acid digestion with concentrated perchloric acid. The released phosphate reacts with ammonium molybdate in the presence of reducing ascorbic acid, forming ammonium phosphomolybdate. This solution, with a blue color, is then measured spectrophotometrically. For this study, both the dried aliquoted lipid extracts and the silica gel containing PL classes separated by HPTLC were placed in glass tubes and treated similarly. After acid digestion at 170°C, the reaction continued with the addition of 1.5% ammonium molybdate (Anedra, Buenos Aires, Argentina), followed by incubation for 5 minutes with 5% ascorbic acid (J.T. Baker, NJ, USA), in a 100°C water bath. Then, the absorbances of samples, blanks, and standards were measured at 800 nm. The measured PL levels were expressed with respect to the dry tissue weight (DTW) of each sample.

#### 7.3 Determination of free cholesterol and cholesterol esters

For separation of neutral lipids, cerebellar tissue lipid extracts were spotted on laboratory-prepared silica gel 60 G plates (Merck, Cat. # 107731, Darmstadt, Germany). While total PL remained at the seeding point, different classes of neutral lipids were resolved using a hexane/ethyl ether/acetic acid solution (80:20:1 v/v). This allowed acceptable separation among free cholesterol (CHOL), diacylglycerols (DAG), free fatty acids (FFA), triacylglycerols (TAG), and cholesterol esters (CE). For band visualization, plates were sprayed with 0.05% 2’,7’-dichlorofluorescein (DCF) (Sigma-Aldrich, D6665) in methanol and inspected under UV light. Lipid standards were co-spotted on plates to allow lipid identification. Free cholesterol and CE were quantified from aliquots taken from the corresponding bands on the preparative TLC plates, after elution from the silica. In both cases, aliquots were dried under N_2_, and cholesterol content was quantified using the Colestat enzymatic AA kit (Wiener Laboratories, Rosario, Argentina). From lipid extracts, aliquots corresponding to 5 µg of phosphorus were dried with N_2_ gas. The dry lipid samples were resuspended in 100 µL of isopropyl alcohol, mixed with 1 mL of the kit reagent, and incubated at room temperature for 20 minutes. The absorbance of the colored product was read at 505 nm in a spectrophotometer. Calibration curves were prepared using a cholesterol standard (2 g/L) provided by the same kit.

#### 7.4 Separation of phospholipids (PL) by classes and determination of composition by HPTLC

To determine the PL composition from cerebellar lipid extracts, commercial 20×10 cm HPTLC plates pre-coated with silica gel (Merck, Cat. # 1056260001), employing two solvent systems, were used. The upper half of the plate was treated with a chloroform/methanol/acetic acid/water (50:37.5:3.5:2 v/v/v/v) mixture to resolve PL (162), while the hexane/ethyl ether (80:20 v/v) mixture was used to separate TAG, TUE, and CE from neutral lipids. After locating the bands on the plates, they were scraped into glass tubes, and their phosphorus content was determined as it was described above (161).

#### 7.5 Determination of lipid-free dry tissue weight (DTW)

The solid residue obtained after lipid extraction from cerebellar tissue was dried in an oven at 110°C for 3 hours and subsequently placed in a desiccator, weighing it multiple times until a constant weight was reached. The resulting lipid-free dry tissue weight (expressed as DTW) represents an estimated weight of the protein and nucleic acid content present in cerebellar samples. This parameter was used to reference and normalize lipidomic data and most calculations described in Section 7, as it eliminates tissue hydration variations as a possible source of error in estimates (163). Statistical comparisons between groups were conducted with Graph Pad Prism 9.0.2 (GraphPad Software Inc.). Data underwent a normality test (Shapiro-Wilk), followed by unpaired Student’s t-test or unpaired Mann-Whitney test (for comparisons between phospholipid classes for which distributions were not normal). Results are graphically represented as mean ± SEM.

### 8 Computational Methods

#### 8.1 General setup

All molecular dynamics (MD) simulations were performed with GROMACS-2018.3 (164–166) patched with Plumed 2.4.3 (167), a convenient plug-in for complex collective variable implementation. The semi-isotropic NPT ensemble at T = 303.15 K (168–170) was used in all production runs, together with a V-rescale thermostat (171) using a 1 ps coupling constant. The pressure was maintained at 1.0 bar with the Berendsen barostat (172), using a compressibility of 3×10 ^ꟷ4^ bar^ꟷ1^ and a 5 ps coupling constant. The fourth-order Particle-Mesh-Ewald method (PME) (173) was used for the long-range electrostatic interactions and a time step of 20 fs was set in all cases. This simulation scheme lets the membrane reach a tensionless equilibrium state after a reasonably long period of equilibration.

The Martini force field (174) with the polarizable water model (175) was applied in all cases along this work, which is widely used for protein-lipid molecular modeling (176–180). As in our previous simulation schemes (181, 182), we used bilayers of 1024 lipid molecules. They had water layers placed both above and below, containing a sufficient amount of water molecules to verify the ample water condition for Martini (>15 CG waters per lipid), which equals to 60 atomistic water molecules per lipid (183).

All protein-membrane simulations were minimized and equilibrated for at least 100 ns to generate properly relaxed initial configurations. Production runs reached the *µ*s scale in all cases. Protein-membrane MD snapshots were created with Visual Molecular Dynamics (VMD) (184) and the academic version of Maestro Molecular Modeling Environment (185). Coarse-grained protein-membrane systems were prepared using the CHARMM-GUI web server (186).

#### 8.2 Coordination

We have used the coordination function as a quantitative measure for protein-protein and protein-lipid interactions (182, 187), using a rational switching function as implemented in Plumed. To quantify the dynamic contacts between different kinds of beads, such as beads that model both amino acids and lipid molecules, it becomes necessary to use a general function as the coordination number (Eq. 1). This analysis focuses on a central bead of interest, and the aim is to evaluate how many beads among a cohort of its neighboring beads are well coordinated with the central one, and how many are not. The coordination function tests each bead-pair (variables i and j) to determine whether that bead-pair can be added to the simple count of well-coordinated bead-pairs, inside a sphere with a user-defined cut-off radius (*r*_0_).

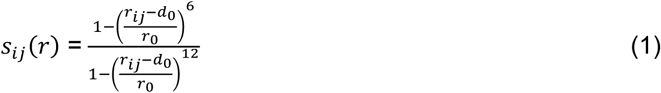

**Equation 1**. *r*_*ij*_ are the three-dimensional distances between beads belonging to different groups (i.e., cholesterol-protein), *r*_0_ is the cut-off distance set to 0.3 nm and parameter *d*_0_ = 0:225 nm corresponds to a mean vdW radius of the included membrane beads species. It follows that *s*_*ij*_ is ≃1 if a contact between beads *i* and *j* is established and *s*_*ij*_ ≃ 0 otherwise. Hence, if any two beads *i* and *j* are separated by a larger distance *r*_*ij*_ ≫ *r*_0_ then the denominator to the 12^*th*^ power makes the total ratio to be ≃ 0. On the other hand, if *r*_*ij*_ ≃ *r*_0_ then both numerator and denominator are comparable making the total ratio to be ≃1.

#### 8.3 Martini beads

In Martini, beads are classified according to the nature of their mutual interactions (polar, non-polar, apolar, charged), and also sub-classified to determine their capability to form hydrogen bonds (donor, acceptor, both, or none) and/or their degree of polarity (from 1 to 5) (174). Cholesterol molecules are modeled by 8 beads, 6 for the sterol and 2 for the tail, with a mainly planar geometry (174). With this mapping of four-to-one coarse-graining, protein-membrane systems, such as the ones included in this study, contain beads that model both amino acids and lipid molecules.

#### 8.4 Identification of CRAC and CARC motifs

We used the human GABA_B2_ subunit sequence (UniProt accession number: O75899) and the Fuzzpro tool from the European Molecular Biology Open Software Suite (EMBOSS) to search for putative transmembrane domain (TM) cholesterol consensus motifs CRAC and CARC. CRAC stands for <u>c</u>holesterol recognition/interaction <u>a</u>mino acid <u>c</u>onsensus and presents a general pattern arrangement as follows: N-terminal-(L/V)-X_1–5_-(Y)-X_1–5_-(K/R)-C-Terminal, where random residues from 1 to 5 can cover the X positions. CARC was named from reading the word CRAC backwards and corresponds to an inverted CRAC motif, N-terminal-(K/R)-X_1–5_-(Y/F/W)-X_1–5_-(L/V)-C-terminal, where once again any 1 to 5 amino acid residues can replace the X position. Fuzzpro search allowed for the retrieval of several compatible patterns for which we selected only not-redundant sequences within the GABA_B2_ TMs. This resulted in the identification of one CRAC (^657^LGIVYAYK^664^) and 3 CARC (^476^KISLPLYSILSAL^488^; ^556^RTWILTV^562^; ^664^KGLLMLFGCFL^674^) matching motifs. We further analyzed coordination over time between cholesterol molecules and each of these sequences.

## Acknowledgments

This work was supported by grants from the CONICET (PIP-13CO01: DM), ANPCyT Argentina (PICT2017-0499 and PICT2021-0314: EMM; PICT2017-1002: DM) and NIH (2 R01 GM083913-41A1: EMM and EV). Also, supercomputing time for this work was provided by CCAD-UNC. The authors would like to thank South Dakota-Biomedical Research Infrastructure Network Proteomics Core Facility (SD-BRIN) (NIH-NIGMS 5P20GM103443-22) for the proteomics analysis, Messrs. Ryan Johnson and Bill Conn from USD-IT Research Computing for their help in the server operation and maintenance, and Raymond D. Astrue for editing the manuscript.

